# SWI/SNF complex-mediated chromatin remodelling in *Candida glabrata* is vital for immune evasion

**DOI:** 10.1101/2023.04.15.536997

**Authors:** Kundan Kumar, Aditi Pareek, Rupinder Kaur

## Abstract

Immune evasion is critical for fungal virulence. However, how the human opportunistic pathogen *Candida glabrata* (*Cg*) accomplishes this is unknown. Here, using micrococcal nuclease-sequencing, RNA-sequencing, macrophage-signalling and genetic analyses, we demonstrate that chromatin reorganization in macrophage-internalized *Cg*, via CgSnf2 (ATPase subunit of the SWI/SNF chromatin remodelling complex), leads to upregulation and downregulation of immunosuppressive seven mannosyltransferase-cluster (*CgMT-C*) and immunostimulatory cell surface adhesin *EPA1* genes, respectively. Consistently, *EPA1* overexpression and *CgMT-C* deletion led to increased IL-1β (pro-inflammatory cytokine) production and reduced *Cg* proliferation in macrophages. Further, *CgSNF2* deletion evoked increased IL-1β secretion, and the consequent killing of macrophage-internalized *Cg*, with elevated IL-1β levels being partially reversed in Akt-, p38-, NF-κB- or NLRP3 inflammasome-inhibited macrophages. Importantly, macrophages respond to multiple *Candida* pathogens via NF-κB-dependent IL-1β production, underscoring NF-κB signalling’s role in fungal diseases. Finally, we present the first genome-wide nucleosome map of macrophage-internalized *Cg* consisting of ∼12,000 dynamic and 70,000 total nucleosomes. Altogether, our findings directly link the nucleosome positioning-based chromatin remodelling to fungal immunomodulatory molecule expression, which dictates *Cg* fate in host immune cells.

## Introduction

Fungal infections cause >1.6 million deaths annually (Bongomin *et al*, 2017). *Candida* species are the most common cause of invasive fungal infections, with *Candida glabrata* (*Cg*) being the second-to-fourth most prevalent *Candida* infectious agent, based upon the geographical location (Chakrabarti, 2015; Bongomin *et al*, 2017; Pfaller *et al*, 2019; Rasheed *et al*, 2020). *Cg* invasive infections are associated with mortality rate exceeding 35% (Wisplinghoff *et al*, 2004; Gupta *et al*, 2015; Meyahnwi *et al*, 2022). *Cg* has recently been classified as a fungal pathogen of high-priority by the World Health Organization (WHO fungal priority pathogens list to guide research, development and public health action).

*Cg* belongs to the *Nakaseomyces* genus, and is also referred as *Nakaseomyces glabrata* (Rasheed *et al*, 2020; Gabaldón *et al*, 2013). *Cg* is evolutionarily closer to the budding yeast *Saccharomyces cerevisiae* (Gabaldón *et al*, 2013; Rasheed *et al*, 2020). *Cg* lacks secreted proteolytic activity and hyphal formation, and predominantly banks upon the stealth-strategy to persist in the host (Kumar *et al*, 2019; Rasheed *et al*, 2020; Kasper *et al*, 2015). *Cg* circumvents the host immune system by inhibiting phagolysosome acidification and suppressing pro-inflammatory cytokine IL-1β production in macrophages (Rasheed *et al*, 2020). Further, the type I interferon signalling contributes to *Cg* immune evasion by dysregulating iron and zinc homeostasis in macrophages (Riedelberger *et al*, 2020b, 2020a).

*Cg* infection to macrophages induces transcriptomic changes in both macrophages and macrophage-internalized *Cg*, with *Cg* relying on multiple mechanisms including chromatin heterochromatinization, carbon metabolism reprogramming, autophagy induction and cell surface reconfiguration to thwart macrophage anti-*Cg* response, and proliferate in macrophages (Kumar *et al*, 2019; Rasheed *et al*, 2020; Kaur *et al*, 2007; Roetzer *et al*, 2010; Seider *et al*, 2011; Kasper *et al*, 2015; Rasheed *et al*, 2018). Notably, *Cg* invokes low levels of IL-1β production in macrophages, and during vaginal and systemic infections (Jacobsen *et al*, 2010; Willems *et al*, 2018; Rasheed *et al*, 2018). Macrophages activate NLRP3 inflammasome and produce IL-1β in spleen tyrosine kinase (Syk) signalling-dependent manner, upon *Cg* infection (Rasheed *et al*, 2018). *Cg* restricts IL-1β secretion in infected-macrophages via a family of eleven putative glycosylphosphatidylinositol (GPI)-linked cell surface-associated proteases (CgYps1-11; CgYapsins) (Rasheed *et al*, 2018). Consistently, *CgYPS1-11* loss led to *Cg* killing in macrophages and clearance from mice organs (Kaur *et al*, 2007; Rasheed *et al*, 2018).

Another important intracellular survival strategy of *Cg* involves chromatin reconfiguration, with CgRtt109 (Histone H3 lysine-56 acetyltransferase) and two subunits of RSC chromatin remodelling complex (CRC), CgRsc3A and CgRsc3B, being required for *Cg* proliferation in macrophages (Rai *et al*, 2012). Macrophage-internalized *Cg* displayed elevated lysine deacetylase activity, and increase and decrease in closed and open-chromatin marks, respectively, with remodelled-chromatin linking metabolic adaptation and energy homeostasis to *Cg* replication (Rai *et al*, 2012). Further, 6 h and 10 h of macrophage internalization rendered *Cg* chromatin resistant to micrococcal nuclease (MNase) digestion, indicating a widespread chromatin reorganization (Rai *et al*, 2012). A nucleosome, consisting of 147 bp DNA wrapped around the histone octamer, is the fundamental repeating subunit of chromatin, and nucleosome dynamics governs chromatin structure and functions (Struhl & Segal, 2013). Nucleosomes occupy defined positions in the genome, with DNA sequence, CRCs, transcriptional regulators and RNA polymerase II (RNAPII) transcription machinery acting in conjunction to maintain nucleosome locations (Struhl & Segal, 2013). Nucleosome positioning control DNA accessibility to various enzymes and regulatory proteins (Struhl & Segal, 2013; Clapier *et al*, 2017). The nucleosome-depleted regions (NDRs), and/or chromatin regions opened up locally by CRCs, facilitate binding of transcription factors and RNAPII machinery to DNA, thereby promoting gene expression (Struhl & Segal, 2013; Clapier *et al*, 2017).

ATP-dependent CRCs mobilize nucleosomes via DNA translocation, histone exchange and/or DNA-histone disengagement (Clapier *et al*, 2017). These belong to four subfamilies: imitation switch (ISWI), chromodomain helicase DNA-binding (CHD), SWI/SNF (SWItch/Sucrose Non-Fermentable) and INO80 (INOsitol requiring), with their catalytic ATPase subunit aiding in DNA translocation (Clapier *et al*, 2017). The SWI/SNF complex in *Cg* and *S. cerevisiae* consists of 12 subunits (Tebbji *et al*, 2017). Transcription factors recruit SWI/SNF complex at specific DNA sites in *S. cerevisiae*, with ScSnf2 involved in DNA binding (Clapier *et al*, 2017). Notably, ScSnf2 localizes to NDRs, and controls transcription initiation by governing +1 nucleosome (transcription start site-associated) position (Nikolov *et al*, 2022).

Despite chromatin architecture being pivotal to intracellular survival of *Cg*, the nucleosome landscape of macrophage-internalized *Cg* is undefined. Here, we report the first genome-wide nucleosome map of macrophage-internalized *Cg* consisting of ∼70,000 nucleosomes, including 12,000 nucleosomes that exhibited changes in position, occupancy or fuzziness. Additionally, we present roles of seven ATP-dependent CRCs in *Cg* pathobiology, underscoring the essentiality of CgSnf2 for immune evasion and virulence. Altogether, our findings unify two central virulence mechanisms, viz., chromatin restructuring and immune evasion, in *Cg*.

## Results

### MNase-sequencing analysis reveals nucleosome dynamics in macrophage-internalized *Cg*

Nucleosome dynamics encompasses nucleosome position shifts, occupancy changes and fuzziness changes (Struhl & Segal, 2013; Chen *et al*, 2013; Clapier *et al*, 2017). To investigate nucleosome dynamics, we performed MNase-seq analysis in 2 h and 10 h macrophage-internalized and RPMI-grown *Cg* cells (Fig 1A). We identified 56,637-71,823 nucleosomes under four-studied conditions (Fig 1B and Appendix Table S1), with 12-24% exhibiting changes in position, occupancy or fuzziness (Fig 1C and D and Appendix Table S2). Most nucleosomes displayed a fuzziness score of 50-60 (Fig EV1A), and most chromosomes contained a high number of dynamic nucleosomes at their ends (Fig 1E). Further, gene bodies displayed the largest number of nucleosomes (Fig EV1B and C). Although overall genomic nucleosome distribution remained unaltered (Appendix Fig S1), 5-6% more dynamic nucleosomes were mapped to intergenic regions in macrophage-internalized *Cg*, compared to RPMI-grown cells (Fig EV1D). Notably, 5% of dynamic nucleosomes were present in promoter regions across the four compared datasets (Fig EV1E), and nucleosome fuzziness was the least observed change in all comparisons (Fig 1D and Fig EV2A-C; and Appendix Table S2).

**Figure 1:**
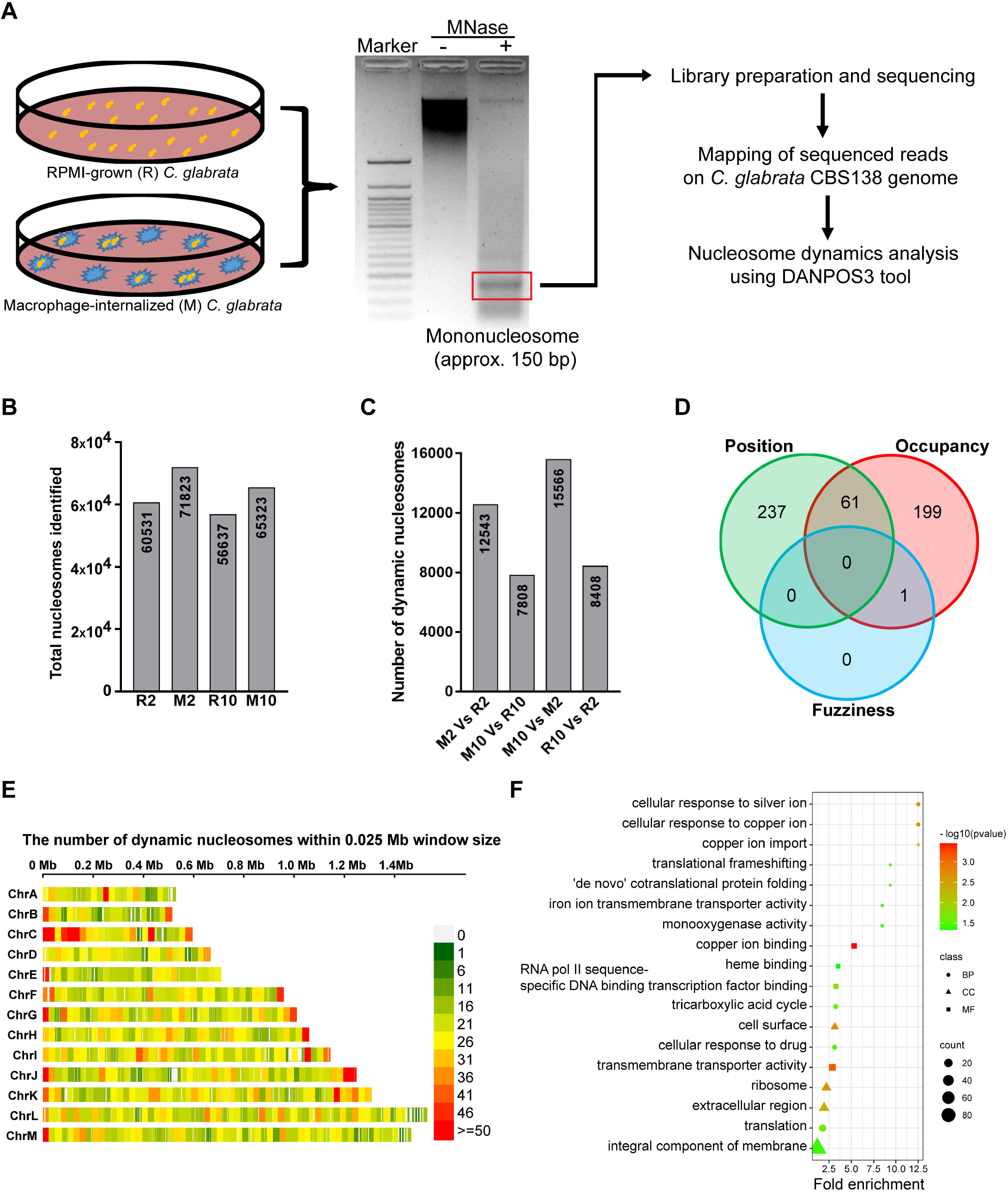
Genome-wide nucleosome dynamics in response to the macrophage milieu. **A,** Experimental design diagram for nucleosome mapping in *Cg wild-type* cells that were grown either in RPMI medium or infected to THP-1 macrophages for 2 h and 10 h. **B,** Total nucleosomes identified using the dpos function of DANPOS3 tool with default parameters. R2: 2 h RPMI-grown, R10: 10 h RPMI-grown, M2: 2 h macrophage-internalized and M10: 10 h macrophage-internalized *Cg* cells. **C,** Dynamic nucleosomes identified in indicated compared datasets. **D,** Venn diagram showing the overlap in the number of genes that exhibited position shift, occupancy and fuzziness changes at their promoter regions in M2 versus R2 comparison. **E,** Chromosome-wise distribution of dynamic nucleosomes in M2 versus R2 comparison. The colour scale indicates nucleosome density. **F,** Gene Ontology (GO) terms for biological process (BP), cellular component (CC) and molecular function (MF) enriched in genes with dynamic nucleosomes at their promoters in M2 versus R2 comparison.

Consistent with macrophage-induced largescale chromatin changes (Rai *et al*, 2012), nucleosomes in 10 h macrophage-internalized *Cg* were more dynamic, compared to 2 h macrophage-internalized *Cg* (Fig 1C), with position shift being the predominant change (Fig EV2B). 10 h macrophage-internalization also led to the highest gain in nucleosome occupancy (Fig EV1F) and the lowest dynamic nucleosome numbers at gene promoters (Fig EV1E and Appendix Table S3), compared to growth in RPMI medium.

David functional enrichment analysis revealed that promoters of the genes, belonging to the GO terms, cell surface, translation, copper ion import and tricarboxylic acid (TCA) cycle, displayed altered nucleosome configurations in 2 h macrophage-internalized *Cg*, with translation gene promoters showing high nucleosome occupancy (Fig. 1F and Appendix Tables S3 and S4). Notably, TCA cycle and translation genes in *Cg* are known to be upregulated and downregulated, respectively, in response to the macrophage milieu (Kaur *et al*, 2007; Rai *et al*, 2012). Further, compared to 2 h, 10 h macrophage internalization led to dynamic nucleosomes in promoters of genes belonging to GO terms, fungal-type cell wall, translation and adhesion of symbiont to host (Fig EV2D and Appendix Table S4). Notably, fungal-type cell wall gene promoters contained dynamic nucleosomes in both 10 h macrophage-internalized and 10 h RPMI-grown *Cg*, compared to 10 h and 2 h RPMI-grown *Cg*, respectively (Fig EV2E and F, and Appendix Table S4), indicating the contribution of chromatin changes to the cell wall gene expression plasticity.

We draw six major conclusions from our MNase-Seq data which report nucleosome dynamics for the first time in a host cell-internalized eukaryotic pathogen. First, promoter regions in *Cg* are nucleosome-depleted, consistent with the previous study (Tsankov *et al*, 2010). Second, 10-20% nucleosomes displayed changes in position, occupancy or fuzziness upon macrophage internalization. Third, macrophage-induced transcriptional downregulation and upregulation, respectively, of translation and TCA cycle genes may be governed by increased and decreased nucleosome occupancy at respective gene promoters. Fourth, nucleosome positioning may govern the expression of cell wall genes in response to the environmental cues. Fifth, compared to gene bodies, dynamic nucleosomes were double in number at promoter and intergenic regions after 2 h macrophage internalization. Finally, position shift was the most common nucleosome dynamic event in *Cg*. Altogether, besides unveiling the macrophage milieu-induced gene-specific nucleosome-pattern in *Cg*, our findings raise the possibility that CRCs, which are essential for nucleosome dynamics, may drive *Cg* survival in macrophages.

### CRCs modulate intracellular survival of *Cg*

*Cg* possesses seven ATP-dependent CRCs. To elucidate their functions, we generated six deletion strains, *Cgsnf2Δ*, *Cgisw1Δ*, *Cgisw2Δ*, *Cgchd1Δ*, *Cgino80Δ* and *Cgswr1Δ*, for non-essential genes that encode putative ATPase subunits of SWI/SNF, ISWI, ISWI, CHD1, INO80 and INO80 CRC subfamilies, respectively (Appendix Fig S2A). Next, we examined growth profiles and virulence-associated traits in generated mutants. Since *Cgrsc3aΔbΔ* has previously been reported to display reduced survival in macrophages and mice (Rai *et al*, 2012), we included this mutant in our analysis as well. Phenotypic profiling revealed pleiotropic stress, and low pH and high iron sensitivity of *Cgsnf2Δ* and *Cgswr1Δ*, respectively, (Fig 2A). *Cgsnf2Δ* grew extremely slowly with a doubling time of 5 h in YPD medium (Appendix Fig S2B and C), and exhibited elongated cell morphology (Appendix Fig S2D). *Cgrsc3aΔbΔ* and *Cgino80Δ* could not utilize alternate carbon sources and displayed reduced growth in iron-surplus medium, with *Cgino80Δ* also exhibiting elevated thermal stress susceptibility (Fig 2A), and 40% longer doubling time than *wild-type* (*wt*) (Appendix Fig S2C).

**Figure 2:**
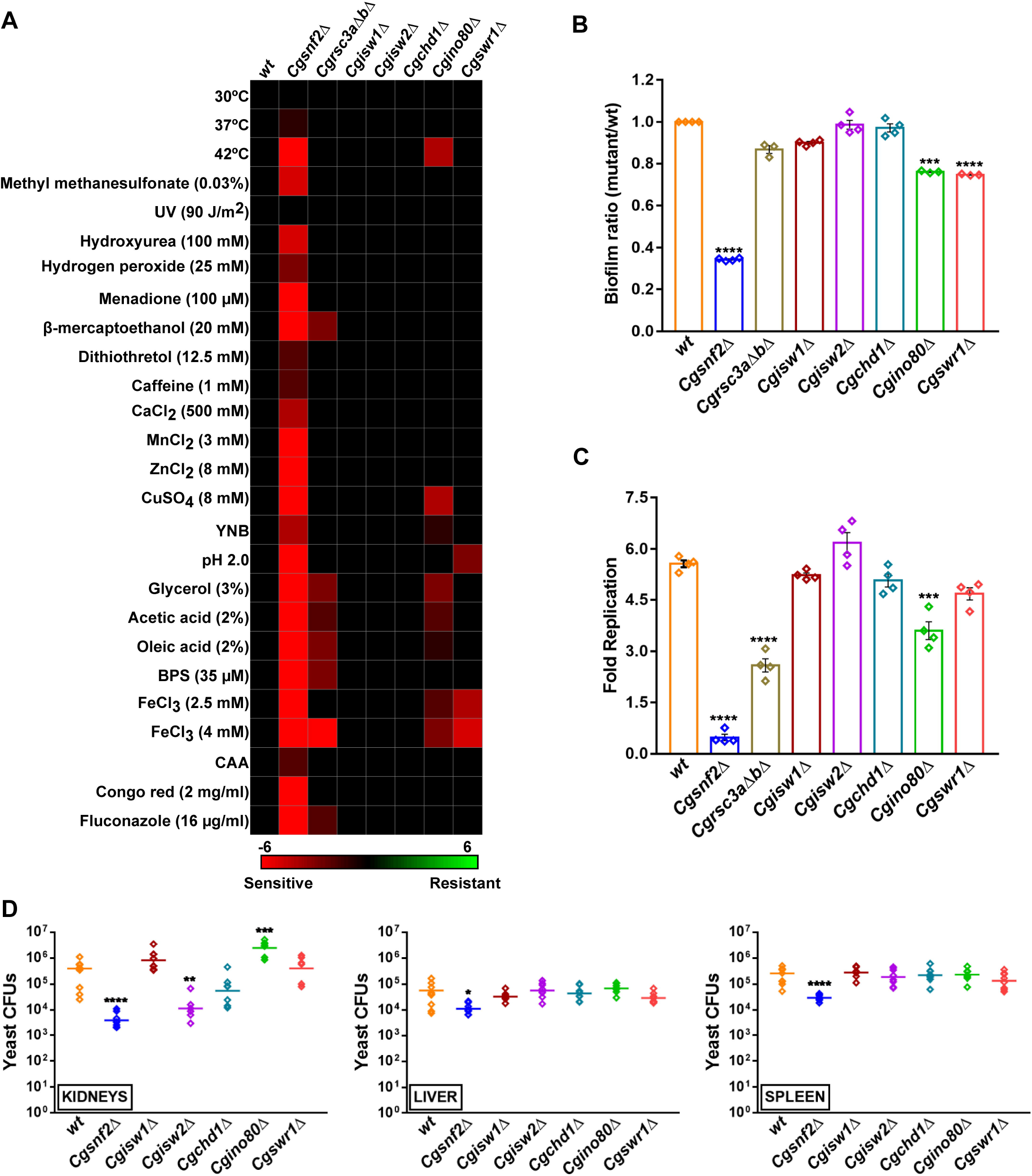
CgSnf2 is essential for intracellular survival and virulence. **A,** Heatmap showing growth profiles of *Cg* strains. The colour code depicts differences between mutant and *wt* growth under the same condition. **B,** Biofilm formation on polystyrene. Data represent mean ± SEM (n = 3-4). Paired two-tailed Student’s t-test. **C,** CFU-based measurement of *Cg* survival in human THP-1 macrophages. Data (mean ± SEM; n =4) represent fold replication at 24 h. Unpaired two-tailed Student’s t-test. **D,** *Cg* survival in the murine model of systemic candidiasis. Diamonds and horizontal line represent fungal CFUs recovered from each mouse, and CFU geometric mean (n = 6-9) for each organ, respectively. Mann-Whitney test.

These distinct stress sensitivity phenotypes notwithstanding, three mutants, *Cgsnf2Δ*, *Cgino80Δ* and *Cgswr1Δ,* were attenuated in their ability to form biofilms on polystyrene surface (Fig 2B). Further, human THP-1 macrophage-infection analysis revealed differential survival of chromatin remodeller-mutants. While *Cgisw1Δ*, *Cgisw2Δ*, *Cgchd1Δ* and *Cgswr1Δ* replicated like *wt* in THP-1 cells, *Cgrsc3aΔbΔ* and *Cgino80Δ* were defective in intracellular proliferation (Fig 2C). Contrarily, *Cgsnf2Δ* was killed in macrophages (Fig 2C), underscoring CgSnf2 essentiality for intracellular survival of *Cg*. Importantly, *Cgsnf2Δ* and *Cgino80Δ* were able to replicate in RPMI medium, albeit slowly than *wt* (Appendix Fig S2E). Further, CgSnf2 was crucial for *Cg* persistence in all three target organs, kidneys, liver and spleen, in the mouse model of disseminated candidiasis, as 5- to 100-fold lower organ fungal load was observed in *Cgsnf2Δ*-infected mice, compared to *wt*-infected mice (Fig 2D). Notably, *Cgisw2Δ*- and *Cgino80Δ*- infected mice displayed 20-fold lower and 7-fold higher renal fungal burden, respectively (Fig 2D), indicating a positive and an antagonistic role for CgIsw2 and CgIno80 in *Cg* fungal load in kidneys, respectively. Altogether, besides implicating ATP-dependent CRCs in diverse stress survival *in vitro*, these data underscore that while CgChd1 and CgIsw1 are dispensable for the examined pathogenesis-associated traits, CgSnf2 is required for *Cg* virulence. Notably, CgSnf2 loss has previously been associated with impaired growth and biofilm formation (Riera *et al*, 2012).

### CgSnf2 suppresses macrophage activation

Among CRC mutants, *Cgsnf2Δ* was most severely compromised for pathogenicity, we, therefore, decided to characterize the *Cgsnf2Δ* mutant further. First, we showed through complementation analysis that the diverse stress susceptibility (Appendix Fig S3A), and elongated cell morphology of *Cgsnf2Δ* (Appendix Fig S3B) is due to the lack of *CgSNF2*. Similarly, *CgSNF2*-expressing *Cgsnf2Δ* displayed *wt*-like survival in macrophages and mice (Appendix Fig S3C and D).

Since *Cg* survival in THP-1 macrophages has earlier been associated with the inhibition of NLRP3 inflammasome- and Syk-dependent IL-1β secretion (Rasheed *et al*, 2018), we next determined if intracellular killing of *Cgsnf2Δ* is owing to elevated IL-1β secretion. We found 15-fold increased IL-1β secretion by *Cgsnf2Δ*-infected THP-1 macrophages, compared to *wt*- infected macrophages (Fig 3A), which was brought down by 4-fold by the NLRP3 inhibitor MCC950 (Fig 3B), with MCC950 also partially rescuing *Cgsnf2Δ*’s intracellular survival defect (Fig 3C). *Cgsnf2Δ* also exhibited 2-fold higher survival in R406 (Syk inhibitor)-treated macrophages, compared to untreated macrophages (Fig 3D). Further, *Cgsnf2Δ* was killed in primary murine macrophages (Fig 3E), with *Cgsnf2Δ*-infected primary macrophages exhibiting 10-fold increased IL-1β levels, and MCC950 treatment diminishing the elevated IL-1β secretion (Fig 3F). These data underscore the conserved response of cultured and primary, as well as, human and mouse macrophages to *Cgsnf2Δ* infection. Altogether, these results implicate CgSnf2 in *Cg*-mediated suppression of the NLRP3 inflammasome activation, and reinforce that impeding IL-1β production is crucial for *Cg* survival in macrophages.

**Figure 3:**
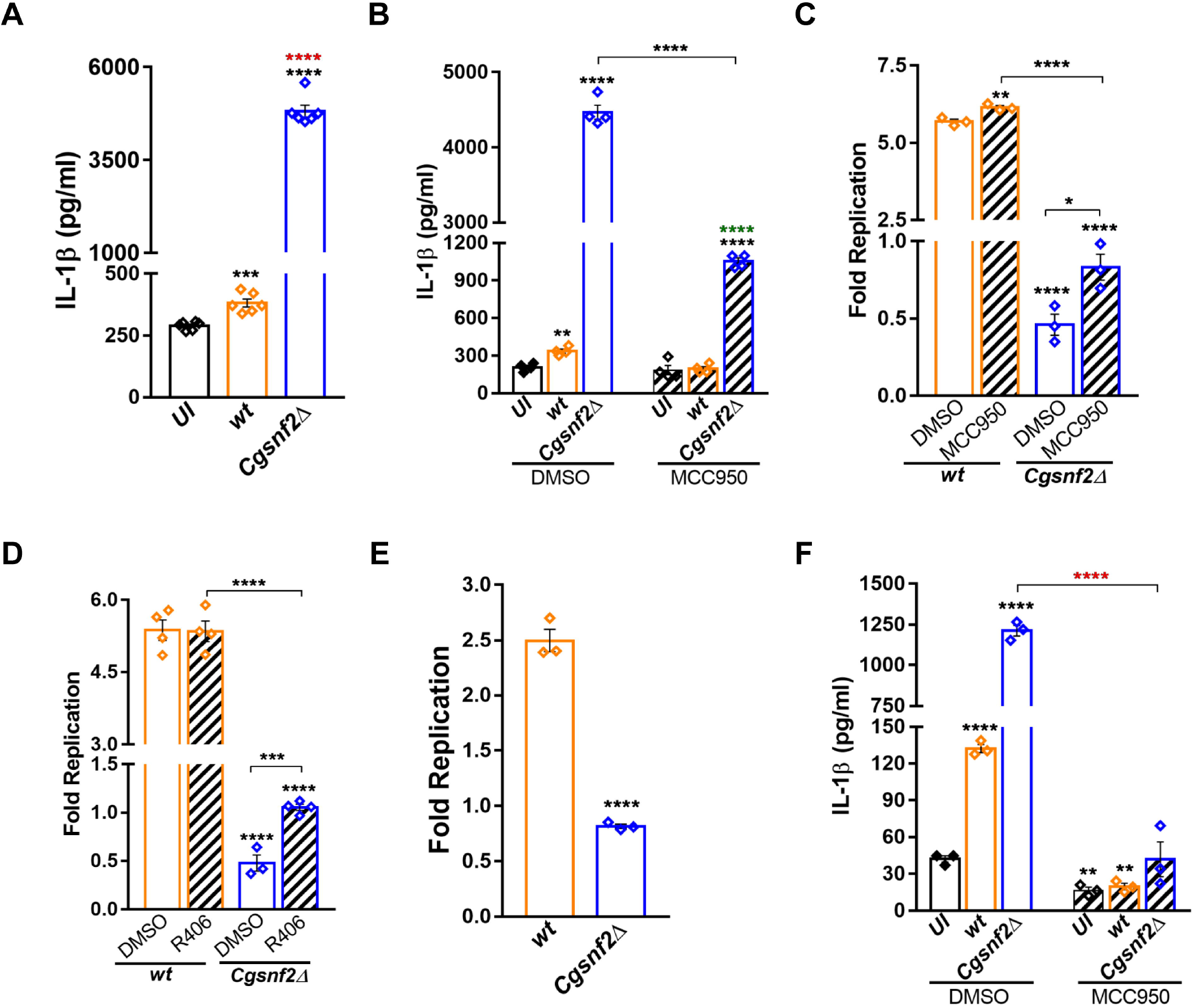
CgSnf2 suppresses host immune cell activation. **A,** Secreted IL-1β levels by uninfected (UI) and *Cg*-infected THP-1 macrophages. Data represent mean ± SEM (n = 6). Unpaired two-tailed Student’s t-test. Black and red asterisks indicate statistically significant differences in IL-1β, compared to UI and *wt-*infected THP-1 cells, respectively. **B,** IL-1β secretion in uninfected or *Cg*-infected THP-1 macrophages that were treated with DMSO (solvent control) or NLRP3 inflammasome inhibitor (MCC950; 15 µM). Data represent mean ± SEM (n = 4). Unpaired two-tailed Student’s t-test. Black and green asterisks represent statistically significant differences in IL-1β, compared to DMSO-treated uninfected and inhibitor-treated uninfected macrophages, respectively. **C and D,** *Cg* survival measurement in DMSO-, 15 µM MCC950 (**C**) or 5 µM R406 (Syk inhibitor) (**D**)-treated THP-1 macrophages. Data (mean ± SEM; n = 3-4) represent fold replication at 24 h. Unpaired two-tailed Student’s t-test. **E,** *Cg* survival measurement in murine peritoneal macrophages. Data (mean ± SEM; n =3) represent fold replication at 24 h. Unpaired two-tailed Student’s t-test. **F,** IL-1β secretion in uninfected or *Cg*-infected murine peritoneal macrophages that were treated with DMSO or 15 µM MCC950. Data represent mean ± SEM (n = 3). Unpaired two-tailed Student’s t-test. Black asterisks represent statistically significant differences in IL-1β, compared to DMSO-treated uninfected macrophages.

### CgSnf2 governs transcriptional response to the macrophage internal milieu

*Cg* persists in macrophages in three-distinct phases, Early, Mid and Late stage, with genome heterochromatinization being a trademark of the Mid-stage (6-12 h of macrophage-ingestion) *Cg* (Rai *et al*, 2012). Notably, CRCs facilitate heterochromatin formation as well as orchestrate transcriptional regulatory proteins-target DNA interactions, that govern gene expression (Clapier *et al*, 2017; Juárez-Reyes & Castaño, 2019). Therefore, to determine the molecular basis underlying CgSnf2-mediated macrophage activation abolishment, we profiled transcriptomes of macrophage-internalized *wt* and *Cgsnf2Δ* at 2 h and 10 h post-infection via RNA-sequencing, and identified differentially expressed genes (DEGs; genes showing ≥ 2-fold change in expression). A total of 1410 (834 upregulated and 576 downregulated) and 192 (162 upregulated and 30 downregulated) genes displayed differential expression in 2 h macrophage-ingested *wt* and *Cgsnf2Δ*, compared to corresponding RPMI-grown cells (Fig 4A and B, and Appendix Table S5). Similarly, 10 h macrophage-internalization led to differential expression of 622 (482 upregulated and 140 downregulated) and 212 (155 upregulated and 57 downregulated) genes in *wt* and *Cgsnf2Δ*, respectively (Appendix Fig S4A-C, and Appendix Table S5), indicating that the magnitude of *Cg* transcriptional response to the macrophage milieu is dependent upon CgSnf2. Consistently, of all DEGs, only 42 genes were found to be CgSnf2-independent (Fig 4C). Of note, compared to *wt*, DEGs in *Cgsnf2Δ* were 7-fold and 3-fold less, upon 2 h and 10 h of macrophage internalization, respectively, indicating that CgSnf2 probably is more important for the initial global response to the varying environment.

**Figure 4:**
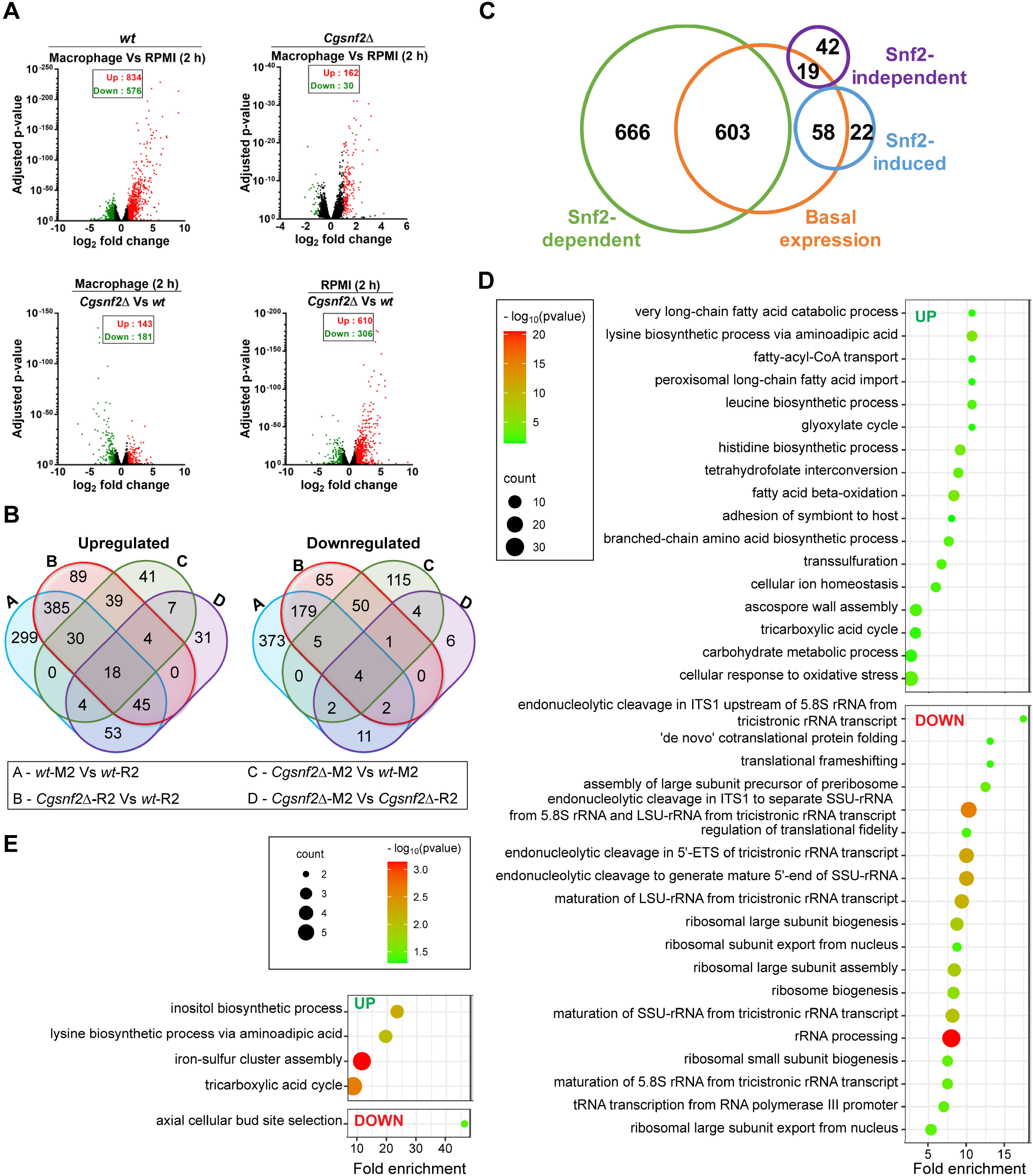
CgSnf2 is required for *Cg* transcriptional response to the macrophage environment. **A,** Volcano plots showing mean fold change of differentially-expressed genes (DEGs) and their adjusted p (false discovery rate-corrected) values. Red, green and black dots indicate upregulated, downregulated and non-DEGs, respectively. **B,** Venn diagrams showing overlap in DEGs among four compared datasets. **C,** Venn diagram illustrating overlap in CgSnf2-regulated genes in 2 h RPMI-grown or 2 h macrophage-internalized *Cg*. Of 1410 DEGs in macrophage-internalized *wt*, expression of 1269 (90%) and 61 (4.3%) genes are dependent and independent of CgSnf2, respectively, with CgSnf2 regulating basal expression of 680 (48%) genes. Additionally, induction of 80 *Cg* genes (5.7%) in macrophages require CgSnf2. **D and E,** Bubble plots showing enriched GO-BP terms for upregulated and downregulated genes in 2 h RPMI-grown *Cgsnf2Δ* **(D)** and 2 h macrophage-internalized *Cgsnf2Δ* **(E)**, compared to 2 h RPMI-grown *wild-type* and 2 h RPMI-grown *Cgsnf2Δ*, respectively.

DAVID functional analysis revealed the enrichment of fatty acid β-oxidation (FAO) and TCA cycle GO terms, in addition to other terms, in upregulated gene sets in 2 h and 10 h macrophage-internalized *wt* (Appendix Fig S4D and Fig EV3 and Appendix Table S6). Contrarily, while rRNA processing, translational frameshifting and ribosome biogenesis terms were enriched in downregulated genes in 2 h macrophage-internalized *wt*, 10 h macrophage internalization led to downregulation of de novo’ IMP biosynthetic process, DNA replication and iron ion homeostasis (Appendix Fig S4D and Fig EV3 and Appendix Table S6). These data are consistent with the earlier microarray-based studies reporting similar gene expression patterns for macrophage-internalized *Cg* (Kaur *et al*, 2007; Rai *et al*, 2012), and underscore reprogramming of the carbon metabolism and shutting down of the translational machinery as characteristic components of *Cg* transcriptional response to the macrophage environment.

Notably, CgSnf2’s role in gene expression is unknown. Our comparison of 2 h RPMI-grown *wt* and *Cgsnf2Δ* transcriptomes revealed that *CgSNF2* deletion led to upregulation of genes involved in FAO, TCA cycle, glyoxylate cycle and adhesion of symbiont to host processes, and downregulation of genes implicated in rRNA processing, translational frameshifting and ribosome biogenesis processes (Fig 4D and Appendix Table S6). Since this transcriptional signature is hallmark of 2 h macrophage-internalized *wt*, it could partly account for the muted-transcriptional response of *Cgsnf2Δ* to macrophage ingestion. Further, the ‘fungal-type cell wall organization’ term was enriched in upregulated genes in 10 h RPMI-grown *Cgsnf2Δ*, compared to the corresponding *wt* (Appendix Fig S4D and Appendix Table S6). We verified the physiological relevance of this observation by performing cell wall component analysis, and found elevated β-glucan and chitin content in *Cgsnf2Δ* (Appendix Fig S4E), which suggests that CgSnf2 aids in cell wall composition maintenance probably by regulating the expression of cell wall organization genes.

Further, *Cgsnf2Δ* responded to 2 h of macrophage-ingestion by upregulating the iron-sulfur cluster assembly, TCA cycle and inositol biosynthetic process, and downregulating the axial cellular bud site selection process (Fig 4E and Appendix Table S6). Notably, TCA cycle and translational frameshifting were upregulated and downregulated, respectively, in 10 h macrophage-internalized *Cgsnf2Δ* (Appendix Fig S4D and Appendix Table S6). These gene expression data collectively suggest that like *wt*, *Cgsnf2Δ* reconfigures carbon metabolism, upon macrophage ingestion, however, it is deficient in closing down the translational machinery, probably because of already-downregulated translation genes.

Notably, contrary to *wt*, *Cgsnf2Δ* showed no activation of the seven mannosyltransferase gene-containing cluster (*CgMT-C*), which included five β-1,2-mannosyltransferase-encoding genes, *CgBMT2-6*, and two α-1,3-mannosyltransferase-encoding genes, *CAGL0B02981g* and *CAGL0B03014g*, after 2 h of macrophage internalization (Fig EV4A-C). Further, comparative transcriptome analysis of 2 h macrophage-ingested *Cgsnf2Δ* and *wt* revealed TCA cycle, gluconeogenesis and FAO downregulation, and NAD transport, adhesion of symbiont to host and fungal-type cell wall organization upregulation (Appendix Fig S4D and Fig EV3 and Appendix Table S6). Since *Cg* adherence to macrophages is the first step in its phagocytosis by macrophages, we next examined how *CgSNF2* loss impacts the expression of cell wall adhesin-encoding genes. Of known 81 adhesins (Xu *et al*, 2020), 21 adhesin genes, including 9 subtelomeric adhesin genes, were differentially expressed in *Cgsnf2Δ* (Fig 5A). Notably, some *EPA* (Epithelial adhesin) genes are subjected to subtelomeric silencing owing to their location at chromosome ends, with CgSnf2 negatively affecting subtelomeric silencing (Riera *et al*, 2012; López-Fuentes *et al*, 2018). CgSnf2 has also been reported to regulate *EPA1* and *EPA6* expression negatively and positively, respectively (Riera *et al*, 2012).

**Figure 5:**
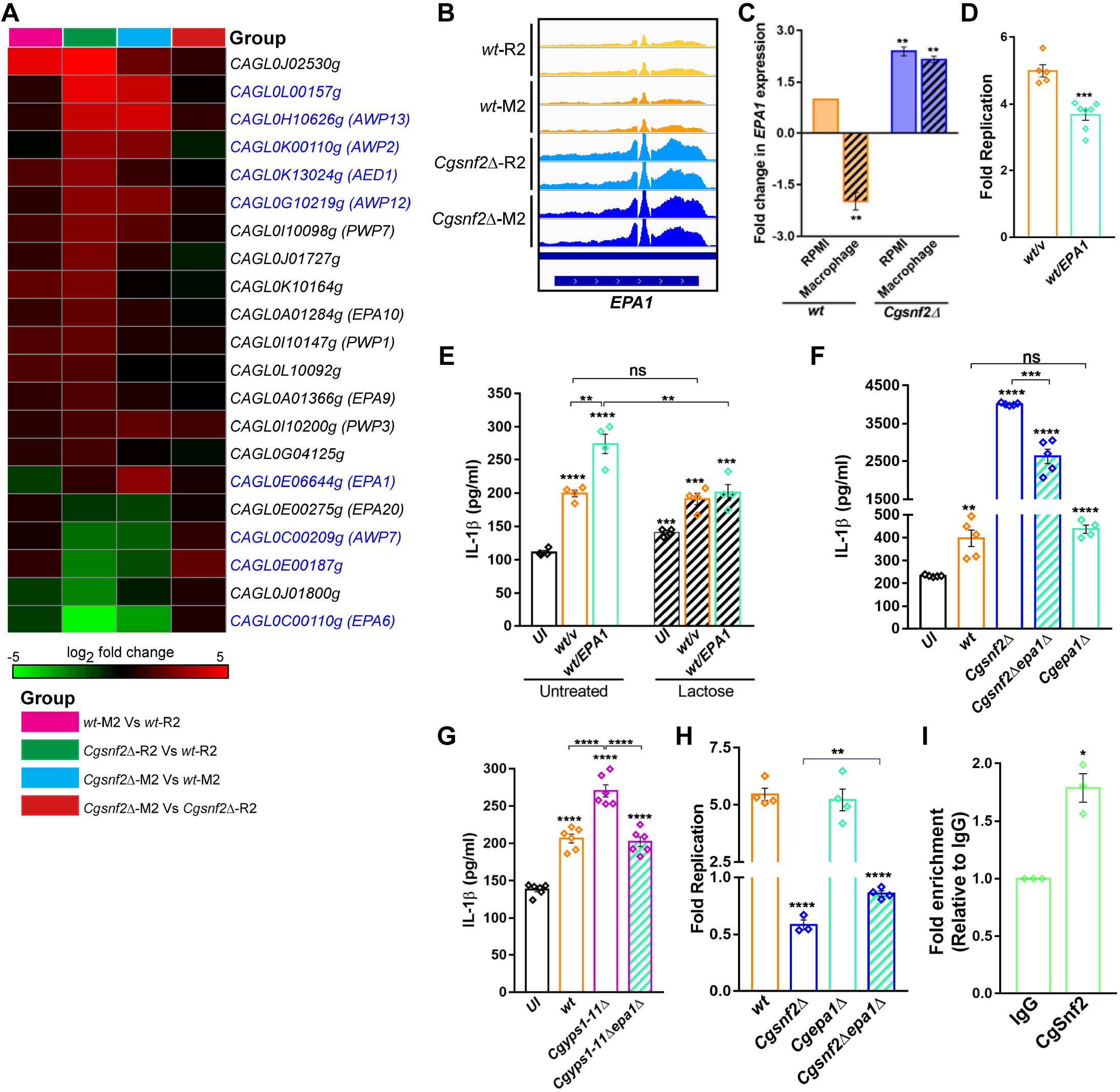
*EPA1* expression is deleterious for *Cg* survival in macrophages. **A,** Heatmap showing differential expression of 21 adhesin genes. Adhesin genes encoded at subtelomeric regions (25 kb from the chromosome ends) are marked in blue colour. R2 and M2 refer to *Cg* grown in RPMI medium and infected to THP-1 macrophages for 2 h, respectively. **B,** Integrative genome viewer (IGV) snapshot of RNA-seq signal at *EPA1* locus (ChrE: 682420 to 685524 bp). All IGV tracks have the same scaling factor [0-750] for the Y-axis. **C,** qPCR-based analysis of *EPA1* expression after 2 h growth in RPMI medium or macrophage internalization. Data mean ± SEM (n = 3) were normalized with *ACT1* mRNA control, and plotted as fold change in gene expression, compared to RPMI-grown *wt* (considered as 1.0). Paired two-tailed Student’s t-test. **D,** *Cg* replication in THP-1 macrophages. Data represent mean ± SEM (n = 5-7). Unpaired two-tailed Student’s t-test. **E,** IL-1β secretion in uninfected (UI) and *Cg*-infected THP-1 macrophages. *Cg* strains were either untreated- or treated with 10 mM lactose, 1 h prior to THP-1 infection, and the infection was continued for 24 h in the presence of lactose. Data represent mean ± SEM (n = 4). Unpaired two-tailed Student’s t-test. **F and G,** IL-1β secretion in uninfected (UI) and *Cg*-infected THP-1 macrophages. Data represent mean ± SEM (n = 4-6). Unpaired two-tailed Student’s t-test. **H,** *Cg* survival in THP-1 macrophages. Data represent mean ± SEM (n = 3-4). Unpaired two-tailed Student’s t-test. **I,** ChIP-qPCR quantification of the level of bound, ectopically expressed SFB-tagged CgSnf2 to *EPA1* promoter. Y-axis label is fold enrichment, with immunoglobulin G (IgG)-control and anti-FLAG (CgSnf2) antibodies. Data represent mean ± SEM (n = 3). Paired two-tailed Student’s t-test.

Based on the *CgMT-C* and adhesin gene expression data, we hypothesized that the perturbed transcriptional regulatory networks in *Cgsnf2Δ* may interfere with the macrophage-induced α- and β-mannan- and adhesin-dependent cell surface remodelling in *Cg*, which may lead to increased IL-1β production and *Cgsnf2Δ* death in macrophages.

### CgSnf2-dependent *EPA1* regulation is pivotal to *Cg*-mediated suppression of IL-1β secretion in macrophages

To test our hypothesis, we first confirmed *CgBMT2* upregulation in macrophage-internalized *wt* by qRT-PCR (Fig EV4D). Next, we generated a mutant lacking all seven genes of the mannosyltransferase cluster, and found that *CgMT-C* deletion led to reduced intracellular proliferation and increased IL-1β secretion (Fig EV4E and F). These results suggest that β-1,2-, and α-1,3-oligomannosides in *Cg* cell wall aid in immunosuppression. Of note, CgBmt2-6 were required for gut colonization in the murine colitis model (Jawhara *et al*, 2012).

Next, to examine the role of adhesins in immunosuppression, we selected *EPA1*, from all differentially-expressed adhesin genes, for further analysis primarily for 3 reasons: (1) Epa1 is the major adhesin for *Cg* adherence to host cells (Cormack *et al*, 1999; Kuhn & Vyas, 2012), (2) *EPA1*-expressing *S. cerevisiae* adhered well to macrophages and stimulated inflammatory cytokine production (Kuhn & Vyas, 2012), and (3) the eleven aspartyl protease-deficient *Cgyps1-11Δ* mutant, with elevated surface-exposed Epa1, was killed in macrophages, due to increased IL-1β secretion (Kaur *et al*, 2007; Rasheed *et al*, 2018). We first verified RNA-seq results by qRT-PCR. *EPA1* transcription was activated and repressed upon *CgSNF2* deletion and macrophage-internalization of *wt* cells, respectively (Fig 5B and C), thereby two possibilities: (1) *EPA1* downregulation aids in suppressing the macrophage pro-inflammatory response, and (2) Increased *EPA1* expression is deleterious to *Cg* survival.

To address these, we performed four experiments. First, we overexpressed *EPA1* from the strong *PDC1* promoter in *wt* and profiled growth in THP-1 cells. We found *wt/EPA1* to display reduced proliferation (Fig 5D), and 3-fold increased IL-1β secretion in macrophages (Fig 5E). Since Epa1 is a calcium-dependent lectin, lactose treatment inhibits its binding to host asialo-lactosyl-containing carbohydrates (Cormack *et al*, 1999). Consistently, THP-1 infection with lactose-treated *wt/EPA1* cells reduced IL-1β secretion significantly (Fig 5E), reinforcing the role of Epa1 in modulating IL-1β production. Second, we deleted *EPA1* gene in the *Cgsnf2Δ* background, and found that *Cgsnf2Δepa1Δ* infection invoked 1.5-fold less IL-1β production in THP-1 macrophages, compared to infection with the single *Cgsnf2Δ* mutant (Fig 5F). IL-1β secretion was similar in response to *wt* and *epa1Δ* infection, probably due to functional redundancy among Epa adhesins (Fig 5F). Third, we performed the same analysis with *Cgyps1-11Δ* after deleting *EPA1*, and found that *Cgyps1-11Δepa1Δ*-infected macrophages secreted 1.3-fold less IL-1β than *Cgyps1-11Δ*-infected cells (Fig 5G). Notably, CgYapsins are required for Epa1 processing from the cell wall, and *Cgyps1-11Δ* contained increased Epa1 levels in its cell wall (Kaur *et al*, 2007). Fourth, we checked the intracellular survival of *epa1Δ* in THP1-macrophages, and found *wt*-like intracellular proliferation, while *Cgsnf2Δepa1Δ* showed 27% better survival than *Cgsnf2Δ* (Fig 5H), highlighting Epa1’s adverse contribution to *Cgsnf2Δ* survival in macrophages. Of note, while the *wt*-like behaviour of *epa1Δ* in THP1-macrophages probably reflects a robust system of functionally-redundant adhesins, the substantial contribution of Epa1 to *Cgsnf2Δ* and *Cgyps1-11Δ* mutants’ phenotypes could be due to deregulated expression/functions of other adhesins in these mutants.

Altogether, these data suggest that Epa1 is immunostimulatory, and acts as a fungal activator of IL-1β induction, and that *EPA1* levels are probably regulated transcriptionally by CgSnf2 via nucleosome repositioning, and post-translationally by CgYapsins through its processing off the cell wall (Kaur *et al*, 2007).

### CgSnf2 binds to *EPA1* promoter

To demonstrate that CgSnf2 directly regulates *EPA1* expression, we performed two experiments. First, through chromatin immunoprecipitation, we showed 2-fold enrichment of CgSnf2 on *EPA1* promoter (Fig 5I), suggesting that CgSnf2 keeps *EPA1* expression in check under normal growth conditions. Second, we mutated the conserved serine (Ser-861) residue in the ATPase domain of CgSnf2 (Appendix Fig S5A and B) to aspartate, and found CgSnf2*^S861D^* to express well (Appendix Fig S5C). However, CgSnf2*^S861D^* could neither rescue stress susceptibility of *Cgsnf2Δ* nor abolish elevated IL-1β secretion in *Cgsnf2Δ*-infected macrophages (Appendix Fig S5D and E), underscoring CgSnf2 catalytic functions, and, by extension, the SWI/SNF complex-mediated chromatin remodelling, in *Cg*-mediated suppression of the host immune response. Consistently, loss of two other subunits of the SWI/SNF complex, CgSnf5 and CgSnf6, led to *Cg* killing and elevated IL-1β secretion in macrophages (Appendix Fig S5F and G). These results unequivocally link the SWI/SNF complex with intracellular survival of, and immune suppression by *Cg*.

### CgSnf2-dependent IL-1β suppression involves NF-κB activation

NLRP3 inhibition in macrophages neither completely reversed IL-1β secretion nor *Cgsnf2Δ* killing (Fig 3), which could be due to other activated immune signalling pathways. Therefore, to define CgSnf2-repressed IL-1β-producing host signalling pathways, we focussed on three pathways, nuclear factor-kappa B (NF-κB), phosphoinositide-3-kinase(PI3K)-protein kinase B/Akt (Akt) and p38 mitogen-activated protein kinase (MAPK) signalling pathways. The NF-κB, a heterodimer of p50 and p65 proteins, is a transcriptional factor which regulates NLRP3 expression and controls the macrophage inflammatory gene expression (Patin *et al*, 2019). Akt and p38 pathways are also implicated in antifungal immunity (Patin *et al*, 2019). We found 1.8-, 2- and 10-fold higher phosphorylation of p65 (regulatory protein of NF-κB), Akt serine/threonine kinase (downstream effector of PI3K signalling), and p38 MAPK in *Cgsnf2Δ*-infected macrophages, compared to uninfected macrophages, respectively (Fig 6A and Fig EV5A and B), indicating a role for CgSnf2 in dampening activation of three other host pathways. Consistently, IL-1β levels were lower in BAY 11-7082 (blocks the inhibitory kappa B protein of NF-κB activation), SH-6 (Akt-specific inhibitor), and SB203580 (p38 inhibitor)-treated *Cgsnf2Δ*-infected macrophages, compared to untreated mutant-infected macrophages (Fig 6B and Fig EV5C and D). Further, *Cgsnf2Δ* infection also resulted in elevated secretion of two other pro-inflammatory cytokines, TNF-α and IL-8, in macrophages, which were partially reversed by all immune signalling inhibitors, but for the NLRP3 inhibitor MCC950 (Fig 6C and D). These data suggest that *C. glabrata* suppresses macrophage activation by controlling the inflammatory response via modulation of the multiple signalling pathways.

**Figure 6:**
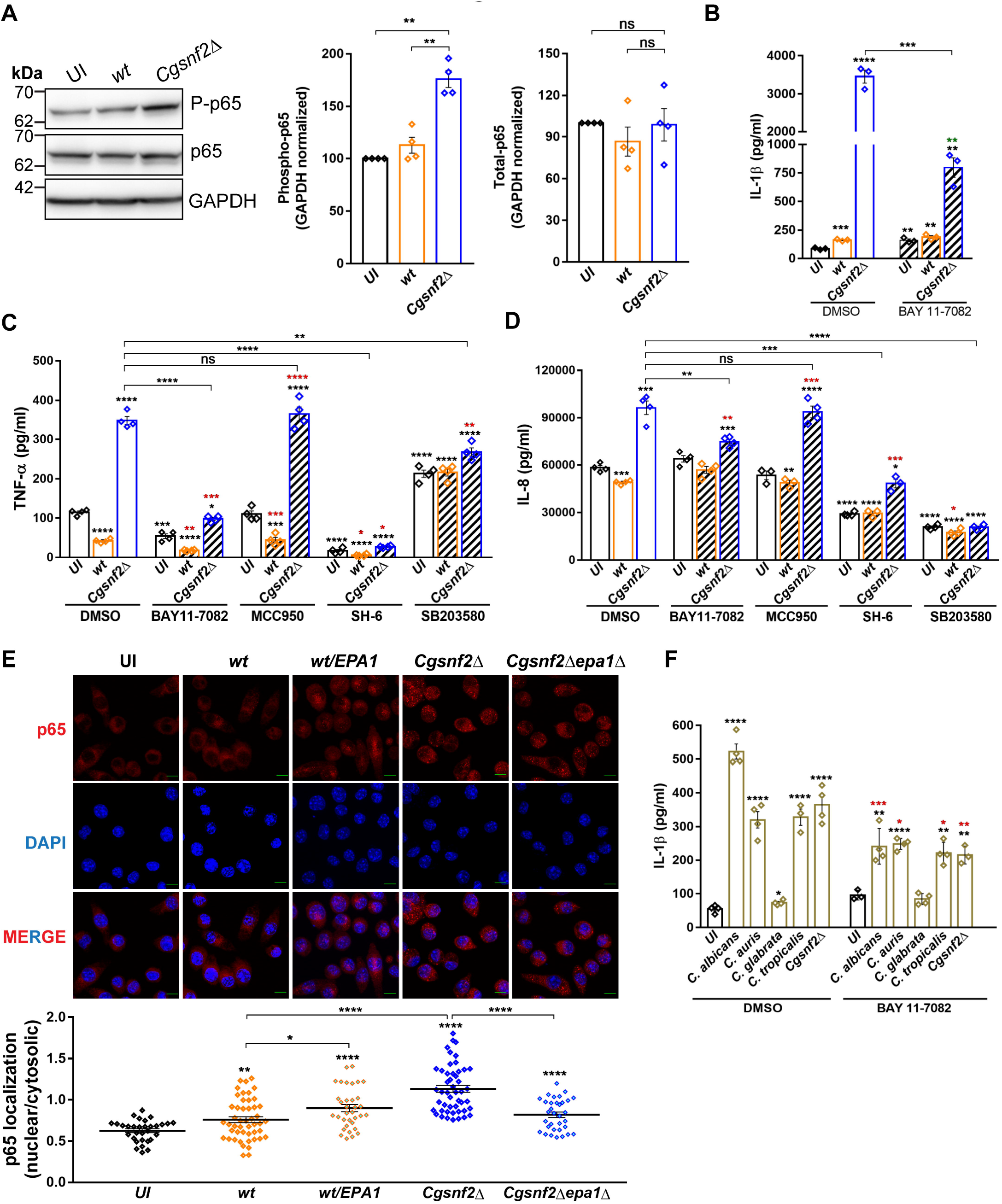
*EPA1* expression governs nuclear localization of NF-κB. **A,** Representative western blots of phosphorylated p65 expression. Bar graphs show fold change in phosphorylation, normalized to GAPDH signal. Data represent mean ± SEM (n = 4). Paired two-tailed Student’s t-test. **B,** IL-1β in uninfected (UI) and *Cg*-infected THP-1 macrophages which were treated with DMSO or NF-κB inhibitor (BAY 11-7082; 10 µM). Unpaired two-tailed Student’s t-test. Black and green asterisks represent statistically significant differences in IL-1β levels, compared to DMSO-treated uninfected and BAY 11-7082-treated uninfected macrophages, respectively. **C and D,** TNF-α **(C)** and IL-8 **(D)** secretion in uninfected (UI) and *Cg*-infected THP-1 macrophages which were treated with DMSO, BAY 11-7082 (10 µM), NLRP3 inhibitor (MCC950; 15 µM), Akt inhibitor (SH-6; 5 µM) or p38 MAPK inhibitor (SB203580; 10 µM). Data represent mean ± SEM (n = 3-4). Unpaired two-tailed Student’s t-test. Black and red asterisks represent statistically significant differences in TNF-α and IL-8 levels, compared to DMSO-treated uninfected and the corresponding inhibitor-treated uninfected macrophages, respectively. **E,** Confocal images showing cellular localization of p65 in uninfected or *Cg*-infected J774A.1 cells, as detected with anti-p65 antibody. DAPI was used to stain macrophage nuclei. Bar = 10 μm. The p65 signal was quantified using ImageJ, and data represent (mean ± SEM; n = 30-50 cells) ratio of nuclear to cytosolic p65. **F,** IL-1β secretion in DMSO or BAY 11-7082 (10 µM)-treated uninfected (UI) THP-1 and THP-1 cells infected with four *Candida* species, *C. albicans*, *C. auris*, *C. glabrata* and *C. tropicalis* at a MoI of 5:1 for 4 h. Data represent mean ± SEM (n = 3-4). Unpaired two-tailed Student’s t-test. Black and red asterisks represent statistically significant differences in IL-1β levels, compared to DMSO-treated uninfected and respective DMSO-treated *Candida spp*-infected THP-1 macrophages, respectively.

Since fungal PAMP (pathogen-associated molecular pattern)-induced signalling pathways may converge on NF-κB activation (Patin *et al*, 2019), which is associated with p65 nuclear translocation, we examined p65 cellular localization in J774A.1 macrophages, as these murine macrophage-like cells have larger cytosol volume. We found 1.5-fold higher translocation of p65 to the nucleus (Fig 6E) in *Cgsnf2Δ*-infected, compared to *wt*-infected macrophages. Further, the role of Epa1 in *Cg*-mediated suppression of NF-κB pathway was demonstrated by enhanced and diminished nuclear translocation of p65 in J774A.1 macrophages infected with *EPA1*-overexpressing *Cg* (Fig 6E) and *Cgsnf2Δepa1Δ* cells (Fig 6E), respectively. Altogether, these data reveal a hitherto unknown facet of *Cg*-mediated host response modulation, via Epa1 adhesin, that involves the master transcriptional regulator, NF-κB, of the inflammatory response.

Finally, to investigate if the NF-κB-dependent inflammatory response represents a pan anti-*Candida* response of macrophages, we infected THP-1 cells with four *Candida* species, *Cg*, *C. albicans*, *C. auris* and *C. tropicalis*. We found increased IL-1β production in all infections, which was reduced upon treatment with BAY 11-7082 (Fig 6F), indicating that NF-κB governs the macrophage inflammatory response against *Candida* pathogens. Notably, *S. cerevisiae* infection of THP-1 cells invoked no IL-1β secretion (Fig EV5E). Altogether, these data indicate that macrophage respond to pathogenic *Candida* yeasts via IL-1β production, and the controlled augmentation of IL-1β has potential for host-directed anti-*Candida* therapy.

## Discussion

*Cg* resides on mucosal surfaces in healthy humans but causes superficial mucosal and life-threatening invasive infections in immunocompromised patients (Fidel *et al*, 1999). Macrophages constitute key host immune defence cells against candidemia, with fungal cell wall structural components, β-glucan and mannan, involved in immune recognition (Erwig & Gow, 2016). Here, we present the first genome-wide nucleosome landscape of macrophage-internalized *Cg*, and establish CgSnf2 as a key determinant of fungal immune evasion. We show that *Cg* modulates NF-κB signalling, by regulating the expression of probable host-recognizable immunostimulatory Epa1 adhesin and immunosuppressive β-1,2-oligomannosides, via CgSnf2, to subvert pro-inflammatory cytokine production in macrophages.

The active immune suppression mechanisms in *Cg* are poorly-understood, and involve impeding phagolysosome acidification and suppressing Syk-dependent IL-1β production (Rai *et al*, 2012; Rasheed *et al*, 2018). The cell wall of *Cg* is the first point-of-contact with the host, and is a multilayered organelle consisting of an inner structural network of chitin, immunostimulatory β-1,3-glucan and β-1,6-glucan, and the outer mannan layer, which is comprised of N-linked or O-linked mannosylated proteins, and is presumed to shield the inner core from immune cells (Rasheed *et al*, 2020; Erwig & Gow, 2016). Dectin-1, Dectin-2, mannose, TLR4, galectin 3 and TLR9 receptors are involved in β-1,3-glucan, α-mannan, mannan and mannoprotein, O-linked mannan, β-mannan, and chitin and DNA recognition, respectively (Erwig & Gow, 2016). Epa1, founding member of the Epa-adhesin family, is a GPI-anchored, glucan cross-linked, calcium-dependent adhesin, possesses a N-terminal PA14 ligand-binding domain, and resides in the outer layer of the cell wall (Cormack *et al*, 1999; Frieman *et al*, 2002; Timmermans *et al*, 2018). Epa1 is shaved off the cell wall in CgYapsins-dependent manner (Kaur *et al*, 2007). Epa1 binds to host glycan ligands containing terminal galactose residue, and mediates adhesion to host epithelial, endothelial and macrophage cells (Cormack *et al*, 1999; Zupancic *et al*, 2008; Kuhn & Vyas, 2012; Timmermans *et al*, 2018). Our data establish Epa1 as a fungal PAMP, whose finely-tuned expression is pivotal to suppress the pro-inflammatory innate immune response. Since Epa1 binding to NKp46 receptor led to *Cg* killing in natural killer cells (Vitenshtein *et al*, 2016), the decrease in surface-exposed Epa1 is likely to constitute a principal *Cg* defence strategy against multiple immune cell-types.

CgSnf2 regulates both basal and macrophage milieu-responsive expression of *EPA1* and other adhesin genes. *Cg* possesses 81 adhesins, with many adhesin genes encoded at subtelomeric regions (Timmermans *et al*, 2018; Xu *et al*, 2020). Nicotinic acid limitation relieves subtelomeric adhesin gene silencing as restricted NAD^+^ availability reduces the activity of the NAD^+^-dependent histone deacetylase Sir2 (Domergue *et al*, 2005; Timmermans *et al*, 2018). Expression of *EPA1*, which is localized 25 kb away from the right telomere of chromosome E, is negatively regulated by the subtelomeric silencing-protein complex, protosilencer Sil2126 and a negative-sequence element (Timmermans *et al*, 2018). We add another regulatory layer of CgSnf2-dependent chromatin remodelling to *EPA1* transcription control which may involve +1 nucleosome position shift, as evidenced in a 60 bp shift towards the *EPA1* gene body in 2 h macrophage-internalized *Cg*, compared to RPMI-grown *Cg* (Appendix Fig S6A). Notably, CgSnf2 regulated the +1 nucleosome position at the promoter of the multidrug transporter gene in *S. cerevisiae* (Nikolov *et al*, 2022). Additionally, we observed nucleosome compaction at the *EPA1* promoter and the 3’ UTR in macrophage-internalized *Cg*, with the average internucleosomal distance being 206 bp and 177 bp in RPMI-grown and macrophage-internalized *Cg*, respectively (Appendix Fig S6B). This complex Epa1 regulation could be pivotal to induce *EPA1* expression for initial infection-stages involving *Cg* adherence to host epithelial and endothelial cells, and repress *EPA1* transcription later to suppress immune recognition and activation. Consistently, early Syk inhibition, following *Cg* ingestion, is pivotal to control *Cg* proliferation effectively in macrophages (Dagher *et al*, 2018), and timely fungal immunogenic molecule-masking and unmasking aids in manipulating immune responses (Cottier *et al*, 2019).

We speculate that in the absence of the hyphal form, *Cg* primarily relies on CgSnf2-dependent dynamic cell surface organization to avoid immune sentinel cells (Fig 7). Accordingly, RNAPII ChIP-seq analysis revealed that *Cg* adhesin genes were upregulated and downregulated, marking immediate (30 min) and late (2 h onwards) response, respectively, to macrophage infection, with CgSnf2 being transcriptionally downregulated at 30 min post-infection (Rai *et al*, 2021). This study also reported 1589 genes to be either constitutively-expressed or temporally-induced in macrophage-internalized *Cg* (Rai *et al*, 2021), which is similar to 1410 DEGs, we identified in 2 h macrophage-ingested *Cg*. Our data suggest that macrophage activation, in response to *Cg* infection, is likely attained by multiple immune signalling pathways. Consistently, loss of the Dectin-1 or Dectin-2 (α-mannan receptor)-mediated Syk signalling individually had no effect on *Cg* replication in macrophages (Dagher *et al*, 2018). Altogether, although our data unequivocally show epigenetic regulation of *Cg* immunomodulatory factors, the nature and distribution of these PAMPs (cell wall-associated or shedded Epa1; α- or β-mannosides), and their immune receptors remain to be identified. Similarly, whether and how Epa1 and mannosides co-operate to antagonize immune-signalling pathways, thereby generating *Cg*-specific immune evasion mechanisms, warrant further studies.

**Figure 7:**
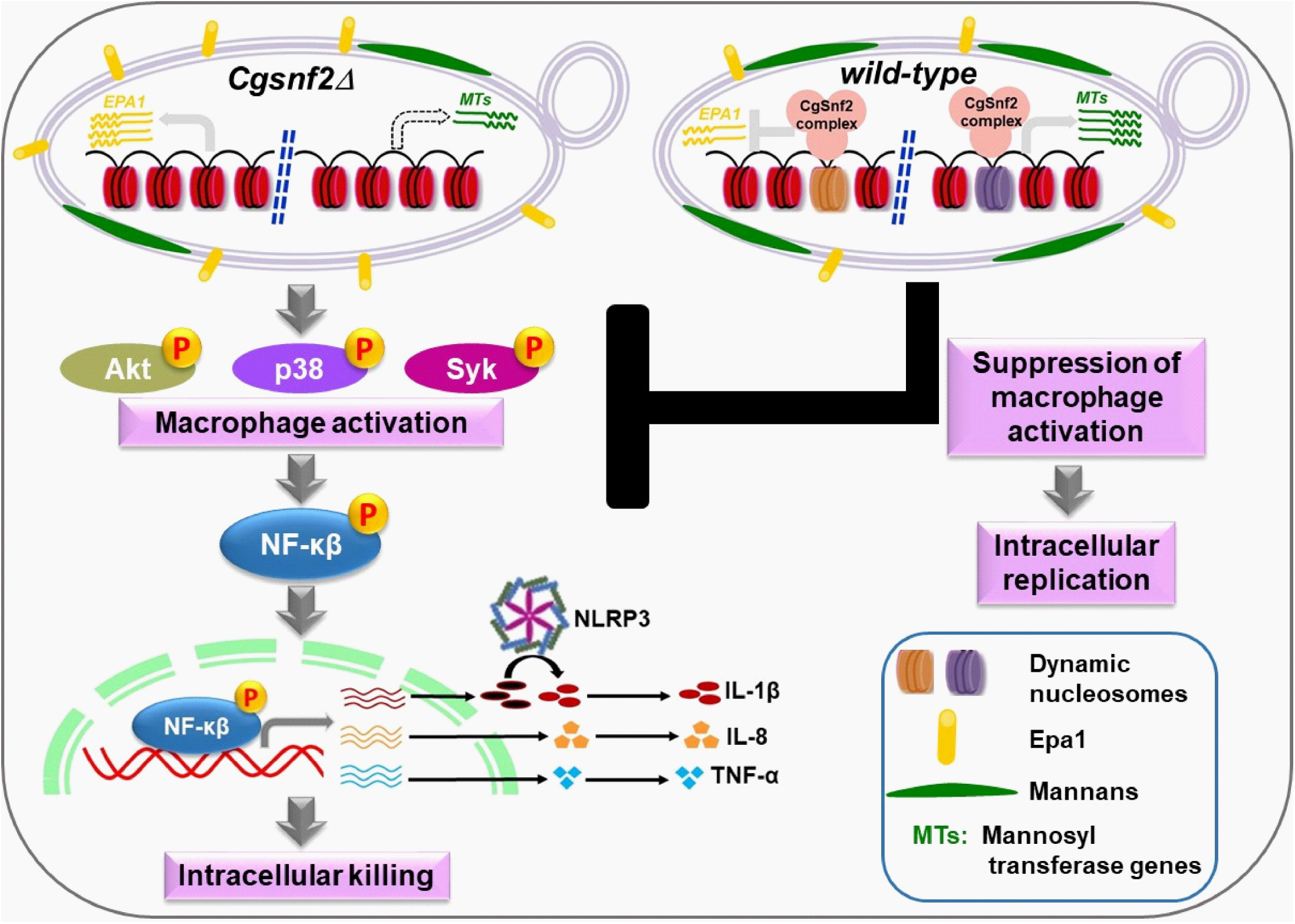
*Cg* uses chromatin remodelling to escape host immune response. A schematic illustrating how CgSnf2 ATPase-mediated chromatin remodelling, during *Cg*-macrophage interaction, results in cell surface reconfiguration, and aids *Cg* suppress signalling pathways in host macrophages.

Microbial pathogens often employ heterochromatin structure to control virulence gene expression (Juárez-Reyes & Castaño, 2019), and nucleosome position changes regulate stress-response genes, with growth and stress-gene promoters exhibiting distinct chromatin features in hemiascomycetous yeasts, including *Cg* (Tsankov *et al*, 2010). *Cg* dynamically switches its chromatin between transcriptionally-active euchromatin and transcriptionally-silent heterochromatin (Rai *et al*, 2012). We mapped 86-102 nucleosomes to the 18.64 kb mannosyl transferase gene-cluster region under R2, M2, R10 and M10 conditions, with 4-22% nucleosomes exhibiting altered occupancy or position (Appendix Tables S1 and S2). Additionally, an increased occupancy at +1 and -2 nucleosomes for *CgBMT5*, and a 90 bp shift towards TSS was observed for *CAGL0B02981*g gene, in 2 h macrophage-internalized *Cg* (Appendix Fig S6C). Notably, the dynamic nucleosome number at *CgMT-C* loci was 3- to 6-fold lower in 10 h macrophage-internalized/10 h RPMI-grown *Cg*, compared to three other datasets (Appendix Table S2). Altogether, these results implicate nucleosome dynamics in controlling *CgMT-C* gene expression, though the significance of observed nucleosome position and occupancy changes remains to be determined. Overall, our data also raise the possibility that the nucleosome pattern on immunomodulatory genes in pathogenic fungi may differ from the gene-averaged nucleosome pattern, which can potentially be useful for therapeutic interventions.

CRCs remodel chromatin by changing composition of, mobilizing or evicting nucleosomes (Clapier *et al*, 2017). *Cg* has seven ATP-dependent CRCs, of which RSC complex ATPase CgSth1 may be essential for *Cg* growth. Since 59% of DEGs in macrophage-internalized *Cg* contained CgRsc3-binding sites (Rai *et al*, 2012), multiple chromatin remodellers may direct nucleosome position. Identifying genomic loci with repositioned/shifted nucleosomes in macrophage-internalized *Cgrsc3aΔbΔ* and *Cgsnf2Δ* mutants, and genome-wide mapping of CgSnf2 and CgSth1, will unveil individualistic CRC contribution. However, given CgSnf2’s essentiality for intracellular survival, its ability to create facile-chromatin regions and recruit specific transcription factors and/or RNAPII machinery may contribute majorly to *Cg* immune evasion.

In conclusion, chromatin reorganization in *Cg* impacts macrophages’ inflammatory response and *Cg* survival. Due to CgSnf2 indispensability for immune evasion, exploring fungal-specific subunits of the SWI/SNF complex (Tebbji *et al*, 2017) as new therapeutic targets holds promise. Additionally, the genome-wide nucleosome map of macrophage-internalized *Cg* will be a useful tool to delineate transcriptional changes arising from chromatin remodelling from those stemming from other gene expression regulatory mechanisms.

## Methods

### Ethics statement

Mice infection experiments were conducted at the Animal House Facility of Centre for DNA Fingerprinting and Diagnostics (CDFD; www.cdfd.org.in) or the CDFD animal facility, VIMTA Labs Limited (http://www.vimta.com), Hyderabad, India in accordance with guidelines of the Committee for the Purpose of Control and Supervision of Experiments on Animals, Government of India. The protocols were approved by the Institutional Animal Ethics Committees of CDFD (EAF/RK/CDFD/22) and VIMTA Labs Ltd (PCD/CDFD/05). Procedures were designed to minimize animal suffering.

### Strains, plasmids and growth conditions

*C. glabrata* (*Cg*) *wild-type* (*wt*) and mutant strains, derivatives of the *Cg*-BG2 strain, were maintained on yeast extract-peptone-dextrose (YPD) medium or minimal yeast nitrogen base medium containing Casamino Acid (CAA) at 30°C. The *Escherichia coli* DH5α strain was used for gene cloning and plasmid propagation, and grown in LB medium containing ampicillin at 37°C. To collect logarithmic (log)-phase *Cg* cells, overnight-grown *Cg* strains were re-inoculated at 0.1 OD_600_ in fresh YPD/CAA medium and cultured for 4-5 h at 30°C.

*Cg* deletion strains were generated, using the homologous recombination-based approach, wherein the *Cg* gene was replaced with the *nat1* gene, that imparts nourseothricin resistance as a selection marker, as described previously (Kumar *et al*, 2020). Despite multiple attempts, *CgSTH1* could not be deleted, which may reflect essentiality of CgSth1 for *Cg* growth. For generation of double mutants, *Cgsnf2Δepa1Δ* and *Cgyps1-11Δepa1Δ*, *CgSNF2* (*CAGL0M04807g*) and *EPA1* (*CAGL0E06644g*) genes were deleted in *Cgsnf2Δ* and *Cgyps1-11Δ* strain backgrounds, respectively. For complementation analysis, *CgSNF2* (5.193 kb) gene along with 5’ (671 bp)- and 3’ (720 bp)-UTR sequences, was cloned at *BamHI* and *XmaI* sites in the pRK1016 (pGRB2.1) vector. For ChIP analysis, *CgSNF2* gene was tagged with the SFB [S-protein(S)-FLAG(F)-Sterptavidin-binding-peptide(B)] epitope at the C-terminus via cloning at Spe*I* and Xma*I* restriction sites in the pRK1349 plasmid, which carries *PDC1* promoter. *CgSNF2^S861D^* was generated by rolling circle method using pRK2404 (*PDC1*- *CgSNF2-SFB*) plasmid DNA as template. For overexpression analysis, *EPA1* (3.105 kb) gene was cloned in *XmaI* and *XhoI* sites, downstream of the strong constitutive *CgPDC1* promoter in the pRK999 vector. The *Candida* strains, plasmids, primers, and antibodies and inhibitors used in this study are listed in and Appendix Tables S7, S8, S9 and S10, respectively.

### Growth analysis

Growth profiles of *Cg* strains in the presence of different stressors were analysed by serial dilution spot or time course assays in solid and liquid medium, respectively. For spot assay, OD_600_ of overnight-grown *Cg* cultures was normalized to 1.0, and cultures were 10-fold serially diluted in phosphate-buffered saline (PBS), followed by spotting 3 μl of each dilution on appropriate medium, and recording growth after 1 to 2 days of incubation at 30°C. For time-course analysis, overnight-YPD medium-grown *Cg* strains were inoculated in fresh YPD medium at an initial OD_600_ of 0.1 and incubated at 30°C with shaking. OD_600_ of cultures was monitored at regular intervals till 36 h, followed by plotting of the absorbance readings as a function of time. Generation time for each strain was calculated during the log-phase (2 h - 8 h) of growth. *Cg* replication in RPMI medium was determined by colony-forming unit-based assay, wherein overnight-YPD medium-grown *Cg* strains were inoculated in RPMI medium containing 10% FBS (fetal bovine serum) at an initial OD_600_ of 0.1 and incubated at 37°C with 5% CO_2_. OD_600_ of cultures was recorded periodically, and appropriate dilution of cultures were plated on YPD medium. After 1-2 days of incubation at 30°C, *Cg* colonies were counted, and CFUs were plotted as a function of time. Mutants showing ≥ 20% longer doubling time than *wt* cells were considered to have retarded growth rate.

### Biofilm formation assay

Log-phase, YPD medium-grown *Cg* cells seeded in a 24-well polystyrene plate at an OD_600_ of 0.5. After 90 min incubation at 37°C, non-adherent *Cg* cells were washed off with PBS, and adherent cells were grown in RPMI 1640 medium containing 10% FBS for 48 h at 37°C, with a medium change at 24 h. After removing unbound cells with PBS washes, adherent *Cg* cells were stained with 0.4% (w/v) crystal violet for 45 min, followed by destaining in 95% ethanol. Absorbance of the destaining solution was recorded at 595 nm, *Cg*-lacking blank well absorbance values were subtracted from *Cg*-containing well absorbance values, and biofilm-forming ability of *Cg* mutants was plotted as the biofilm ratio which represents the mutant/*wt* absorbance values. Mutants showing ≥ 1.25-fold reduction in biofilm ratio were considered to be attenuated for their biofilm-forming capacity.

### Mice infection analysis

For animal infection studies, overnight-grown *Cg* strains in YPD medium at 30^ο^C were collected, washed twice with sterile PBS and suspended in PBS to a final cell density of 20 OD_600_. 100 μl of cell suspension was injected into the tail vein of 6-8 week-old female BALB/c mice. At 7^th^ day post-infection, mice were euthanized using CO_2_ and three organs (kidneys, liver and spleen) were excised out. The harvested organs were homogenized in PBS, and homogenates were appropriately diluted and plated on YPD medium supplemented with penicillin and streptomycin antibiotics. After 1-2 days of incubation at 30°C, *Cg* colonies were counted. The fungal burden in each organ was calculated by multiplying the colony number with the appropriate dilution factor.

### *Cg*-macrophage interaction analysis

Human THP-1 monocytic cells (1 x 10^6^ cells in each well of a 24-well tissue culture plate) were treated with phorbol 12-myristate 13-acetate (16 nM) for 12 h for differentiation into macrophages at 37°C with 5% CO_2_, followed by 12 h cell recovery in RPMI-10% FBS medium. THP-1 macrophages were infected with overnight, YPD medium-grown *Cg* cells at a multiplicity of infection (MoI) of 0.1. Non-phagocytosed *Cg* cells were removed after 2 h, and the infection was continued for another 22 h. At 2 h and 24 h post-infection, extracellular *Cg* cells were removed with PBS, THP-1 cells were lysed in water and lysates were plated on YPD medium. After 2 days of growth at 30°C, *Cg* colonies were counted and the fold replication of *Cg* in THP-1 cells was calculated by dividing 24 h CFUs by 2 h CFU counts. % phagocytosis was determined by dividing 2 h CFUs by 0 h CFUs (Number of *Cg* cells infected to THP-1 macrophages). Mutants showing ≥ 2-fold change in fold replication, as compared to *wt* cells, were considered to have altered survival in macrophages.

Secreted cytokines were measured by enzyme-linked immunosorbent assay (ELISA). For this, THP-1 macrophages were infected with *Cg* at a MoI of 1.0 for 24 h, as described above, and culture supernatants were collected and centrifuged at 1000 rpm to remove any cell debris. IL-1β, IL-8 and TNF-α in cleared supernatants were measured using the ELISA kits following manufacturer’s protocol. For inhibitor treatment, PMA-treated THP-1 macrophages were pre-treated with inhibitors or DMSO solvent for 2 h, prior to *Cg* infection, and *Cg*-THP1 co-culture was carried out in the presence of inhibitor or DMSO for 24 h.

For isolation of primary macrophages, 6-8-week-old BALB/c mice were intraperitoneally injected with 1.5 ml of 4% (w/v) thioglycollate. After 5 days, mice were euthanized, and macrophages were collected in DMEM from the mouse peritoneal cavity. Macrophages were centrifuged, suspended in DMEM medium containing 10% heat-inactivated FBS and were seeded in a 24-well plate. *Cg* survival and secreted IL-1β in primary mouse peritoneal macrophages were measured in a similar way, as that for THP-1 cells.

### Nucleosome dynamics analysis by MNase-Seq

For MNase-seq, PMA-activated THP-1 (2.2 x 10^7^) cells were infected with overnight YPD medium-grown *wt* cells at a MOI of 1:1, and macrophage-internalized *Cg* cells were collected post 2 h and 10 h infection, after lysing infected macrophages in ice-cold water. Macrophage lysates were gently vortexed to separate *Cg* cells form macrophage debris, followed by centrifugation at 6000 rpm for 5 min at 4°C. The compact *Cg* cell pellet was collected and suspended in ice-cold water, followed by centrifugation at 10,000 rpm for 5 min at 4°C. This step was repeated 3-4 times to obtain *Cg* cell pellet that was devoid of macrophage debris. As a control, overnight YPD medium-grown *wt* cells were grown in RPMI-10% FBS medium for 2 h and 10 h, and cell pellets were collected. After PBS washes, 2 h and 10 h RPMI-grown and macrophage-internalized *wt* cell pellets were suspended in MNase-digestion buffer [10 mM Tris-Cl (pH 8.0) and 1 mM CaCl_2_], and lysed using glass beads. After lysate centrifugation at 13000 rpm at 4°C for 10 min, lysates (1 mg) were incubated with MNase (10 units/1 mg lysate; NEB #M0247S) at 37°C for 60 min. The MNase digestion was stopped by adding Stop Buffer (8.6% SDS and 0.007 M EDTA), followed by digestion with proteinase K (20 mg/ml) first at room temperature for 30 min, and later at 65°C for overnight. DNA was isolated with PCI (Phenol:Chloroform:Isoamyl alcohol) extraction, precipitated with sodium acetate and ethanol, and was suspended in nuclease-free water, followed by RNase A digestion for 30 min at 37°C. DNA was run on 2% agarose gel, and bands of ∼ 150 bp, corresponding to mononucleosomal DNA fragments, were excised, purified using QIAquick extraction kit, and were sent to Scigenom, Kochi, India (http://www.scigenom.com/) for library preparation and sequencing. The concentration and quality of purified mononucleosomal DNA was determined by Qubit, Nanodrop and Agilent Tapestation analysis, followed by library generation using the NEBNext Ultra II DNA Library Prep Kit (Illumina), following the manufacturer’s instructions. The library quality was assessed using the Agilent Bioanalyzer High Sensitivity DNA chip.

The prepared libraries were sequenced (2×100 bp paired-end sequencing) on the HiSeq 2500 (Illumina) platform, and 40-60 million, high-quality 100 bp reads, with 93 % of the total reads passing ≥30 phread score, were generated for each sample. Data were analysed by DeepSeeq Bioinformatics, Bengaluru, India (https://www.deepseeq.com/). The adapter sequences and low-quality reads were removed using Trimmomatic (version 0.39). Trimmed sequence reads were aligned to the *C. glabrata* genome version s04-m01-r06 (www.candidagenome.org) using bowtie2, version 2.5.0. The percentage alignment varied between RPMI-grown (89-97%) and macrophage-internalized (6-26%) samples. The SAM files generated by bowtie2 were converted to BAM format, sorted and indexed using Samtools, version 1.7. The DANPOS3 software(Chen *et al*, 2013) was used to identify the occupancy, position and fuzziness of nucleosomes. For this, clonal reads from BAM files were first removed, followed by quantile normalization. Next, individual dpos, dpeak, and dregion functions from DANPOS3 were used to identify positions, peaks, and regions occupied by nucleosomes in individual samples as well as for compared datasets. Annotation of the called nucleosome position was done using gene start and stop coordinates for 5604 genes, obtained from the C_glabrata_CBS138_version_current_features.gff file version S04-m01-r06 downloaded from http://www.candidagenome.org/download/gff/C_glabrata_CBS138/, by employing the closest function from the Bedtools software version v2.28.0. The –flank_up and –flank_down parameters were set to 500 and 1000, respectively, for TSS (Transcription Start Site) -based analysis, and to 500 and 500, respectively, for TTS (Transcription Termination Site)-based analysis. Nucleosomes were identified in about 5300 genes in all four-studied conditions. Lastly, three filters, Position shift: ≥ 50 bp shift, Occupancy change: ≥ 2-fold change (FDR ≤ 0.05) or Fuzziness change: ≥ 1.5-fold change (FDR ≤ 0.05), were applied to identify dynamic nucleosomes.

### Immunoblotting analysis

For signalling pathway analysis, PMA-activated macrophages were either left uninfected or infected with *wt* and *Cgsnf2Δ* cells at a MOI of 1:1. After 4 h infection, macrophages were washed with ice-cold PBS, scrapped and centrifuged at 2000 rpm for 5 min. Cell pellets were suspended in freshly-prepared NETN lysis buffer [250 mM NaCl, 5 mM EDTA (pH 8.0), 50 mM Tris-Cl (pH 8.0), and 0.5% Nonidet P-40] containing protease (cOmplete Mini, Roche) and phosphatase (PhosSTOP, Roche) inhibitors, and incubated for 30 min at 4^ο^C. Cell lysates were sonicated for 15 cycles (30 sec ON/30 sec OFF), with the Diagenode bioruptor sonicator and centrifuged at 13000 rpm for 10 min. 120 µg proteins were resolved on 10% SDS-PAGE, and probed with different antibodies. For quantification, intensities of the bands of interest were quantified from independent immunoblots using the ImageJ densitometry software, and normalized with respective GAPDH signal intensities. Values were plotted as a bar graph, considering the signal in control samples as 1.0. For *Cg* protein analysis, log-phase *Cg* cells were lysed using the glass beads, and lysates (80 μg) were resolved on 10% SDS-PAGE, followed by probing with anti-FLAG antibody.

### *Cg* cell wall analysis

For quantification of major cell wall components, overnight YPD medium-grown *wt* and *Cgsnf2Δ* mutant strains were inoculated in fresh YPD medium at 0.1 OD_600_ and incubated at 30^ο^C for 4 h. Log-phase cells (2.0 OD_600_) were collected, washed and suspended in PBS. Cells were either left untreated or stained with calcofluor white (2.5 µg/ml), fluorescein isothiocyanate-labeled concanavalin A (1 µg/ml) or aniline blue (12.5 µg/ml) for 15 min at room temperature for measurement of chitin, mannan and β-glucan, respectively. After PBS washes, fluorescence intensity of ∼50,000 cells was recorded on BD FACS ARIA III flow cytometer at an excitation wavelength of 355 nm and an emission wavelength of 433 nm. All data were analysed with the FlowJo software, and the background fluorescence was corrected by subtracting the mean intensity fluorescence value of unstained samples from that of respective stained samples. Data were presented as the mean intensity ratio which indicates *Cgsnf2Δ*/*wt* fluorescence intensity values.

### RNA-sequencing analysis

2 h and 10 h RPMI-medium and macrophage-internalized *wt* and *Cgsnf2Δ* cells were harvested and total RNA was extracted using RNeasy kit (Qiagen), followed by removal of DNA contamination, if any, using DNase I. The purified total RNA was sent to National Genomics Core (NGC) at CDFD, Hyderabad (http://ngc.cdfd.org.in/) for sequencing, which involved mRNA enrichment using Poly(A) mRNA Magnetic Isolation Module, library preparation using NEB Ultra II Directional RNA Library Prep Kit and 150 bp paired-end sequencing on the Illumina NextSeq 2000 platform. Sequenced reads were processed and mapped on to the *C. glabrata* CBS138 reference genome using HISAT 2.1.0 aligner.

For gene expression quantification, the counts of mapped reads for each gene were considered using the Feature counts tool, followed by sequencing depth normalization of raw read counts using the DESeq2 tool. Differentially expressed genes were classified based on two criteria: ≥ 2-fold change in expression and adjusted p-value of ≤ 0.05.

### Imaging analysis

For morphology analysis, log-phase *Cg* cells were visualized using the LSM700 confocal microscope equipped with 63X/1.44 NA objective. For cell wall chitin staining, log-phase *Cg* cells were stained with calcofluor white (1.25 µg/ml) for 30 min at room temperature. After PBS washes, cells were imaged using LSM700 confocal microscope equipped with 63X/1.44 NA objective. All images were processed using the ZEN blue software.

For immunofluorescence analysis, J774.1 murine macrophage-like cells (1 x 10^6^) were seeded on a coverslip, and infected with *wt* and *Cgsnf2Δ* cells at a MoI of 1:1. At 2 h post-infection, un-phagocytosed *Cg* cells were removed, followed by 10% FBS-containing DMEM medium addition and incubation for 2 h at 37°C, 5% CO_2_. As a control, uninfected J774A.1 cells were grown under similar conditions. J774. 1 cells were washed with ice-cold PBS and fixed with 3.7% (v/v) formaldehyde for 15 min. After fixation, cells were permeabilized with 0.5% (v/v) Triton X-100 in PBS for 5 min, washed twice with PBS and blocked with 10% (v/v) FBS in PBS for 1 h at room temperature. Cells were probed overnight with anti-NF-κB p65 antibody at 4°C, followed by incubation with AlexaFluor 568-conjugated secondary antibody for 1 h. Coverslips were washed, air-dried and mounted in the VECTASHIELD anti-fade mounting medium containing DAPI. Cells were visualized, and Z-stack images were acquired throughout the nucleus at 1 μm interval using the Leica confocal microscope (63X/1.44 NA objective), and processed using the LAS X software. Each maximum-intensity projection (MIP) image was constructed using 8-12 confocal image subsets.

### Quantitative RT-PCR (qRT-PCR) analysis

RNA was isolated from 2 h and 10 h RPMI-grown and macrophage-internalized *Cg* cells using the RNeasy kit (Qiagen), followed by DNase I digestion. 1 µg DNase I-treated total RNA was used for cDNA synthesis using the Superscript III reverse transcriptase, DyNAmo ColorFlash SYBR Green qPCR kit was used to perform qRT-PCR, and gene expression was determined by the 2^-ΔΔCT^ method. *GAPDH/ACT1* expression was used as an internal reference control to normalize the gene expression data.

### Chromatin immunoprecipitation (ChIP) analysis

ChIP was performed, as described previously (Bhakt *et al*, 2022). Briefly, *CgSNF2-SFB*-expressing *Cgsnf2Δ* was grown in CAA medium for 4 h, followed by formaldehyde crosslinking. Next, cells were lysed in buffer containing 1 mM EDTA [pH 8.0], 50 mM HEPES [pH 7.5], 0.1% sodium deoxycholate [w/v], 140 mM NaCl, 1× protease inhibitor cocktail, and 1% Triton X-100 by bead-beating, and sonicated for 40 min [30 s pulses of on and off] to obtain 250-500 bp small chromatin fragments. A fraction of the clear supernatant was used as Input fraction, while the remainder was subjected to immunoprecipitation with anti-IgG or anti-FLAG antibody. After de-crosslinking, DNA was purified using phenol:chloroform:isoamyl alcohol, and used as template for qRT-PCR. Results are presented as enrichment over IgG.

### Functional enrichment and statistical analysis

The David tool (https://david.ncifcrf.gov/) with default settings was used for enrichment of gene ontology (GO) terms for biological process, cellular component and molecular function, in genes with dynamic nucleosomes at their promoters and in differentially-expressed genes. Terms with the q-value of ≤ 0.05 were considered as enriched in gene datasets. Statistical significance was determined using the two-tailed Student t-test or the nonparametric Mann-Whitney test in the GraphPad Prism software. Error bars indicate standard error of the mean (s.e.m). The multiple amino acid sequence alignment was done using the Clustal W tool. Asterisks were used to represent p-values: * p ≤ 0.05, ** p ≤ 0.01; *** p ≤ 0.005; **** p ≤ 0.001. The p value of ≤ 0.05 was considered significant.

## Data availability

The raw MNase-seq and RNA-seq data will be deposited to the NCBI’s Gene Expression Omnibus (GEO).

## Acknowledgements

We would like to thank Dr. Pranjali Pore and staff of the Experimental Animal Facility of CDFD for their help with mice experiments. We are indebted to Vandana Sharma and Romila Moirangthem for her help with MNase-Seq experiments and *C. glabrata* deletion strain generation, respectively.

## Author contributions

KK conceived the study. KK and RK designed the study. KK and AP performed experiments and acquired data. KK, AP and RK analysed data. KK prepared tables and figures. KK and RK wrote the manuscript with input from AP.

## Supplementary information

This manuscript contains Expanded View Figures (EV1 to EV5).

This manuscript contains Appendix data (Appendix Figures S1 to S6, and Appendix Tables S1 to S10).

## Expanded View Figures (EV1 to EV5)

### Expanded View Figure legends

**Figure EV1 (related to Fig 1):**
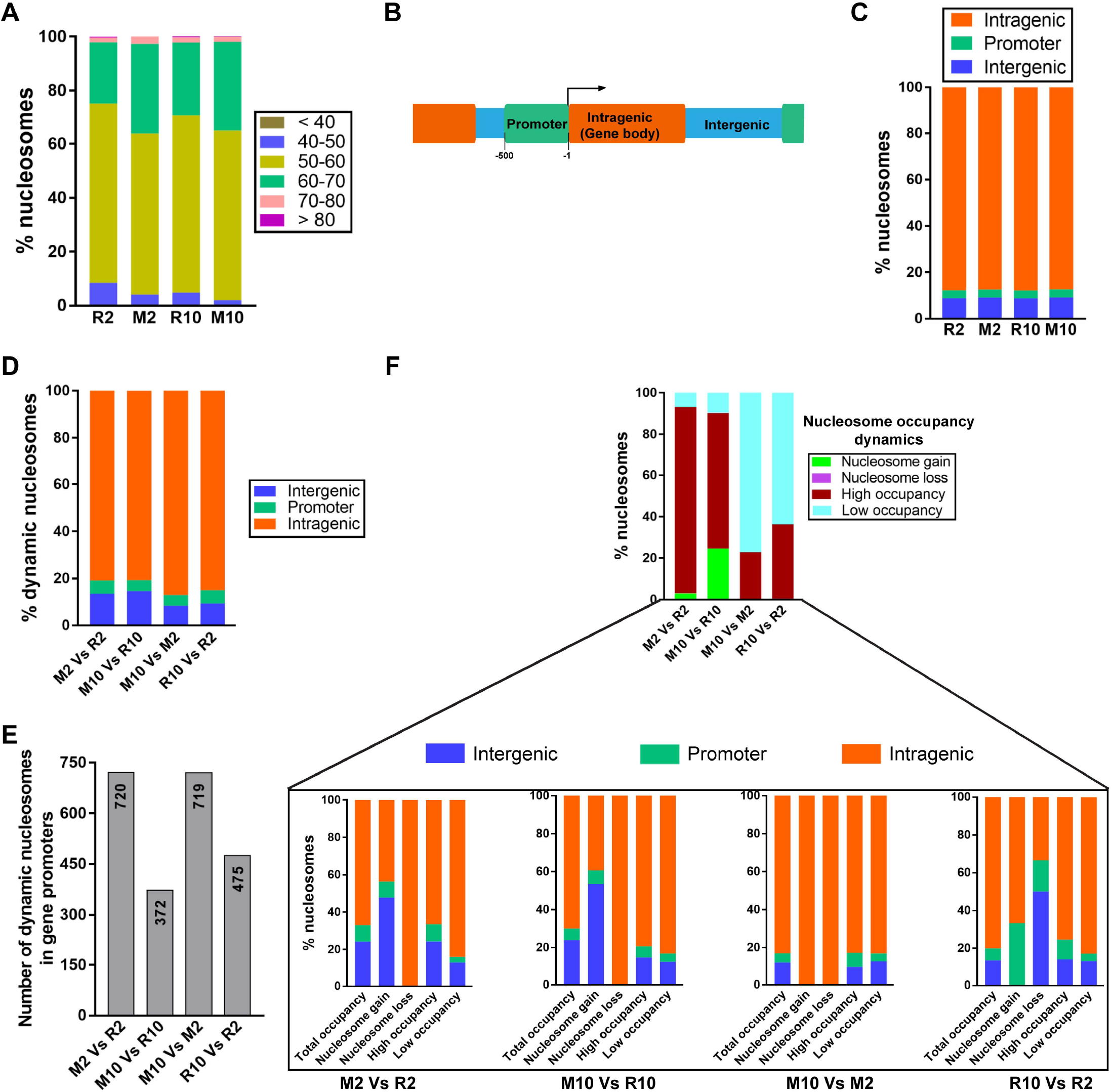
Nucleosome dynamics in macrophage-internalized *Cg*. **A,** Percentage of fuzzy nucleosomes. R2: 2 h RPMI-grown, R10: 10 h RPMI-grown, M2: 2 h macrophage-internalized and M10: 10 h macrophage-internalized *Cg*. **B,** A schematic of a genomic locus divided into three regions for nucleosome mapping: Promoter [0-500 bp upstream of the Transcription Start Site (TSS)], intragenic (gene-encoding sequence) and intergenic (region between the stop codon of one gene and the promoter or the stop codon of the next gene). The arrow indicates the direction of transcription. **C,** Nucleosome distribution across three genomic regions. **D,** Dynamic nucleosomes across three genomic regions. **E,** Dynamic nucleosomes in gene promoters. The exact number is indicated in the upper edge of the bar. **F,** Percentage of four types of nucleosome occupancy dynamics across three genomic regions.

**Figure EV2 (related to Fig 1):**
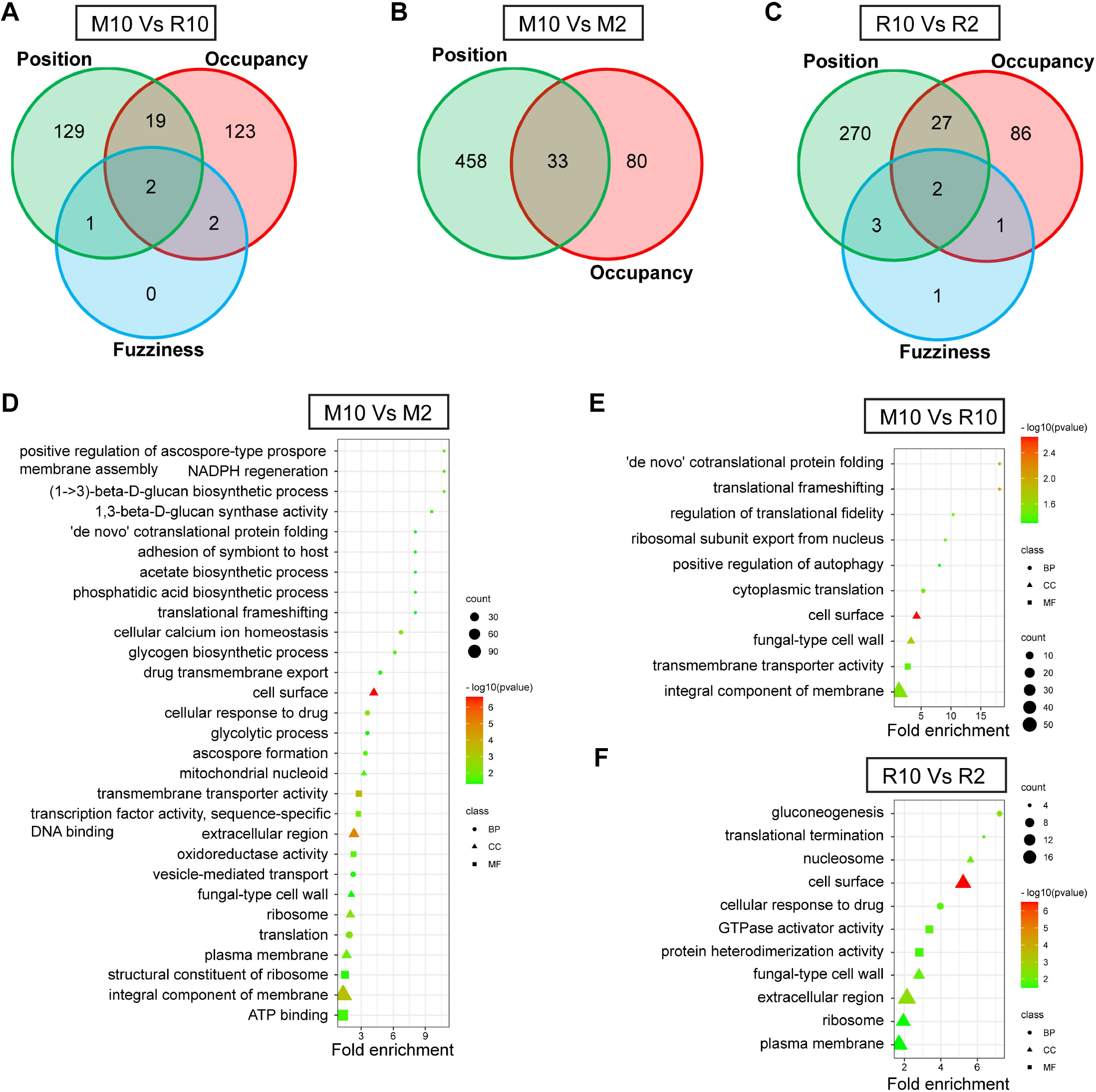
Genes involved in host adhesion contain dynamic nucleosomes at their promoters. **A-C,** Venn diagrams illustrating overlap in the number of genes which showed nucleosome position shift, occupancy and fuzziness changes at their promoters in M10 versus R10 **(A)**, M10 versus M2 **(B)** and R10 versus R2 **(C)** comparison. **D-F,** GO terms for biological process (BP), cellular component (CC) and molecular function (MF) enriched in genes with dynamic nucleosomes at their promoters in M10 versus M2 **(D)**, M10 versus R10 **(E)** and R10 versus R2 **(F)** comparison.

**Figure EV3 (related to Fig 4):**
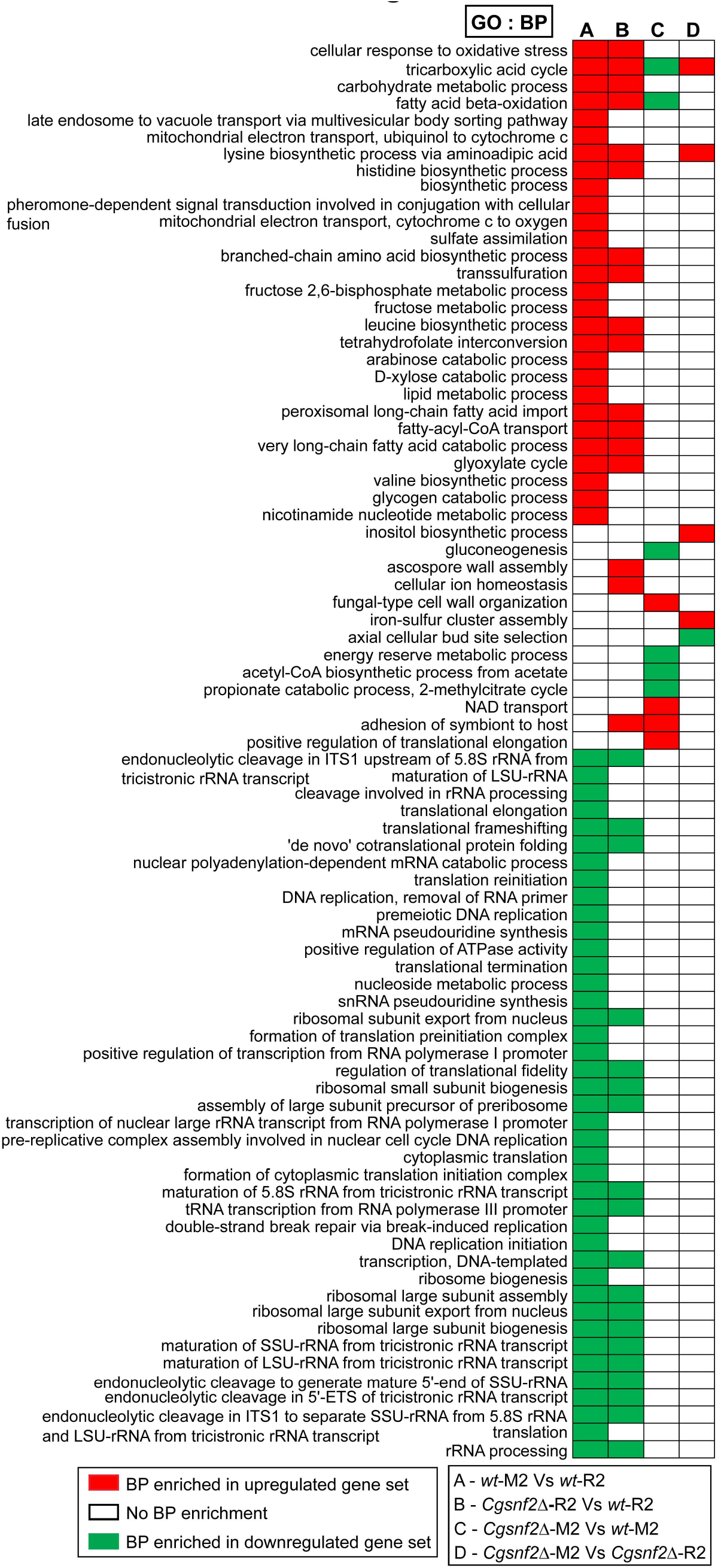
Heatmap showing enriched Gene Ontology-Biological Process (GO-BP) terms in four compared datasets.

**Figure EV4 (related to Fig 5):**
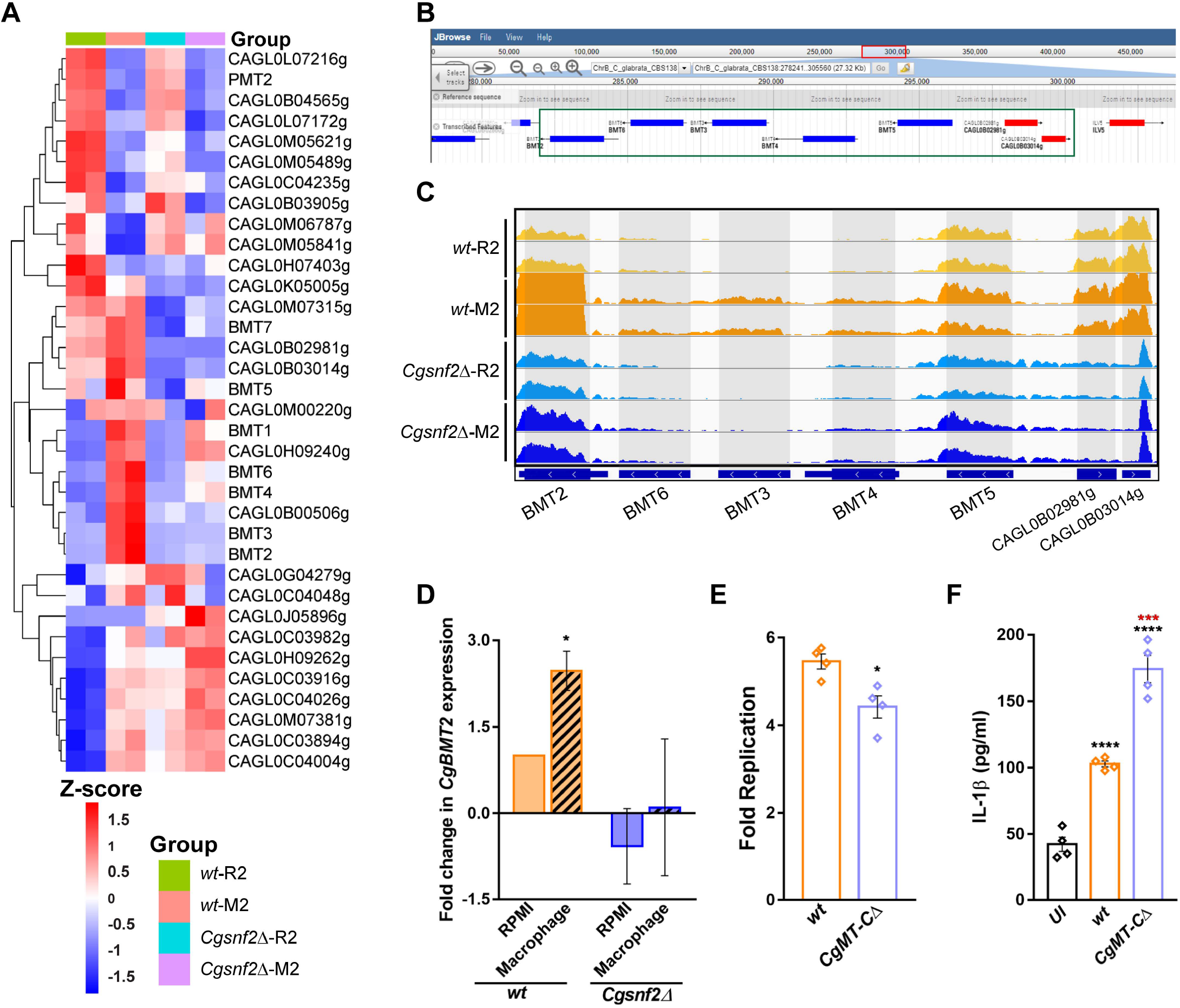
CgSnf2-dependent activation of a 7-mannosyltransferase gene cluster is pivotal to supress host activation. **A,** Heatmap showing normalized read counts (presented as z-score) of 35 mannosyltransferase (MT)-encoding genes in 2 h RPMI-grown (R2) and macrophage-internalized (M2) *wt* and *Cgsnf2Δ*. **B,** JBrowse snapshot (www.candidagenome.org) of the seven (five-β and two-α) mannosyltransferase genes present in tandem on the Chromosome B. **C**, Integrative genome viewer (IGV) snapshot of RNA-seq signal at *MT* loci (ChrB: 282186 to 300029 bp) in *wild-type* and *Cgsnf2Δ* grown either in RPMI-medium (R2) or incubated with THP-1 macrophages (M2) for 2 h. All IGV tracks have the same scaling factor [0-500] for the Y-axis. **D,** qPCR-based analysis of *CgBMT2* expression in 2 h macrophage-internalized *wt* and *Cgsnf2Δ*. Data (mean ± SEM; n = 3) were normalized with *ACT1* mRNA control, and represent fold change in gene expression, compared to RPMI-grown *wt* (considered as 1). Paired two-tailed Student’s t-test. E, *Cg* intracellular replication analysis. The *CgMT-CΔ* lacks ∼19 kb genomic region containing seven mannosyltransferase genes. Data represent mean ± SEM (n = 4). Unpaired two-tailed Student’s t-test. **F,** IL-1β secretion in THP-1 macrophages. Data represent mean ± SEM (n = 4). Unpaired two-tailed Student’s t-test. Black and red asterisks represent statistically significant differences in IL-1β compared to uninfected (UI) and *wild-type*-infected (*wt*) macrophages, respectively.

**Figure EV5 (related to Fig 6):**
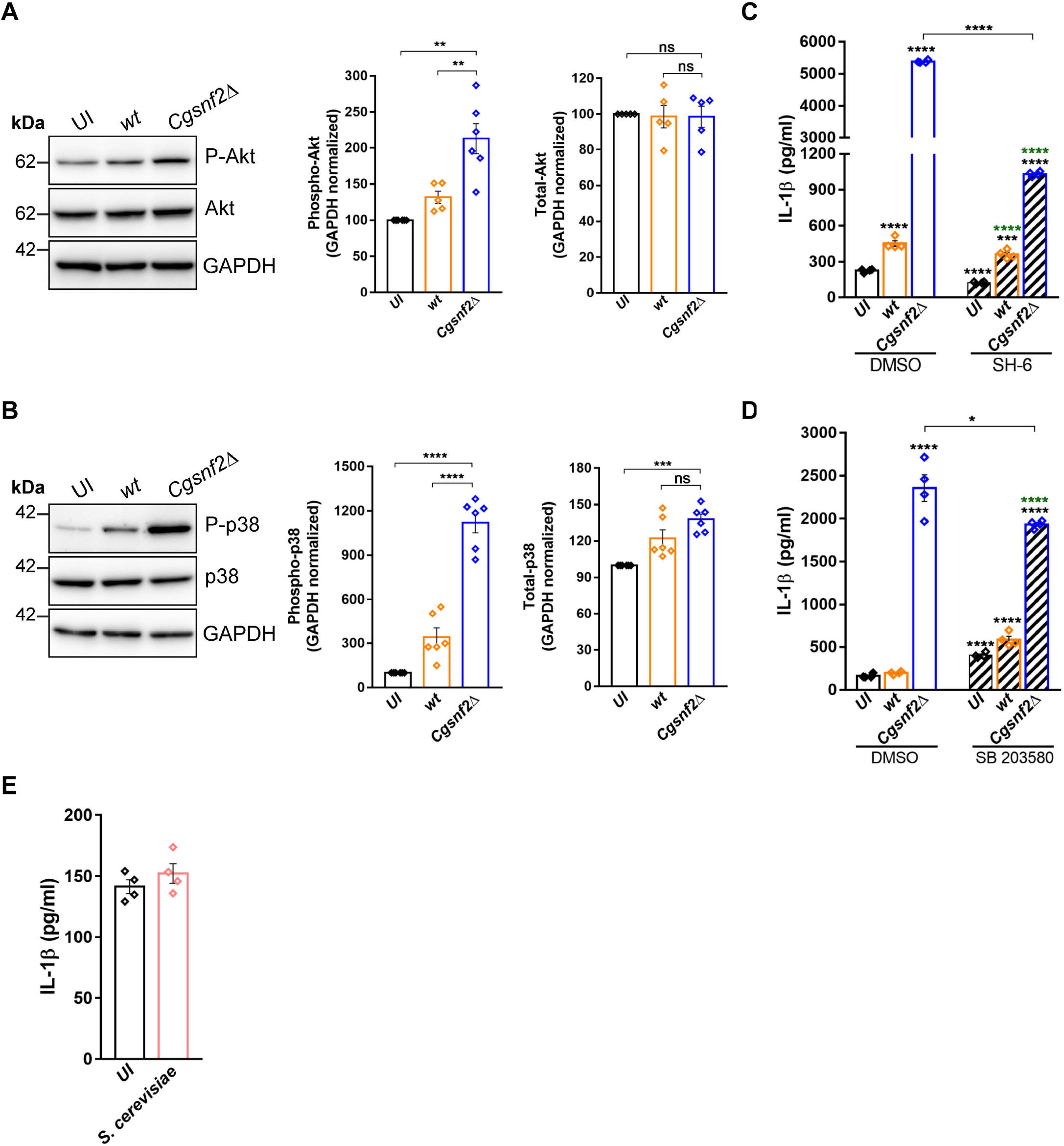
CgSnf2 supresses Akt and p38 activation. **A and B,** Representative western blots of phosphorylated Akt (**A**) and p38 (**B**) expression. Bar graphs show fold change in phosphorylation, normalized to GAPDH signal. Data represent mean ± SEM (n = 5-6). Paired two-tailed Student’s t-test. **C and D,** IL-1β secretion in uninfected and *Cg*-infected THP-1 macrophages which were treated with DMSO or Akt inhibitor (SH-6; 5 µM) (**C**; n=4) and p38 inhibitor (SB203580; 10 µM) (**D**; n=4). Unpaired two-tailed Student’s t-test. Black and green asterisks represent statistically significant differences in IL-1β levels, compared to DMSO-treated uninfected and inhibitor-treated uninfected macrophages, respectively. **E,** IL-1β secretion in uninfected and *S. cerevisiae*-infected THP-1 macrophages. Data represent mean ±SEM (n = 4).

## Appendix data (Appendix Figures S1 to S6, and Appendix Tables S1 to S10)

### Appendix Figures legends

**Appendix Figure S1:**
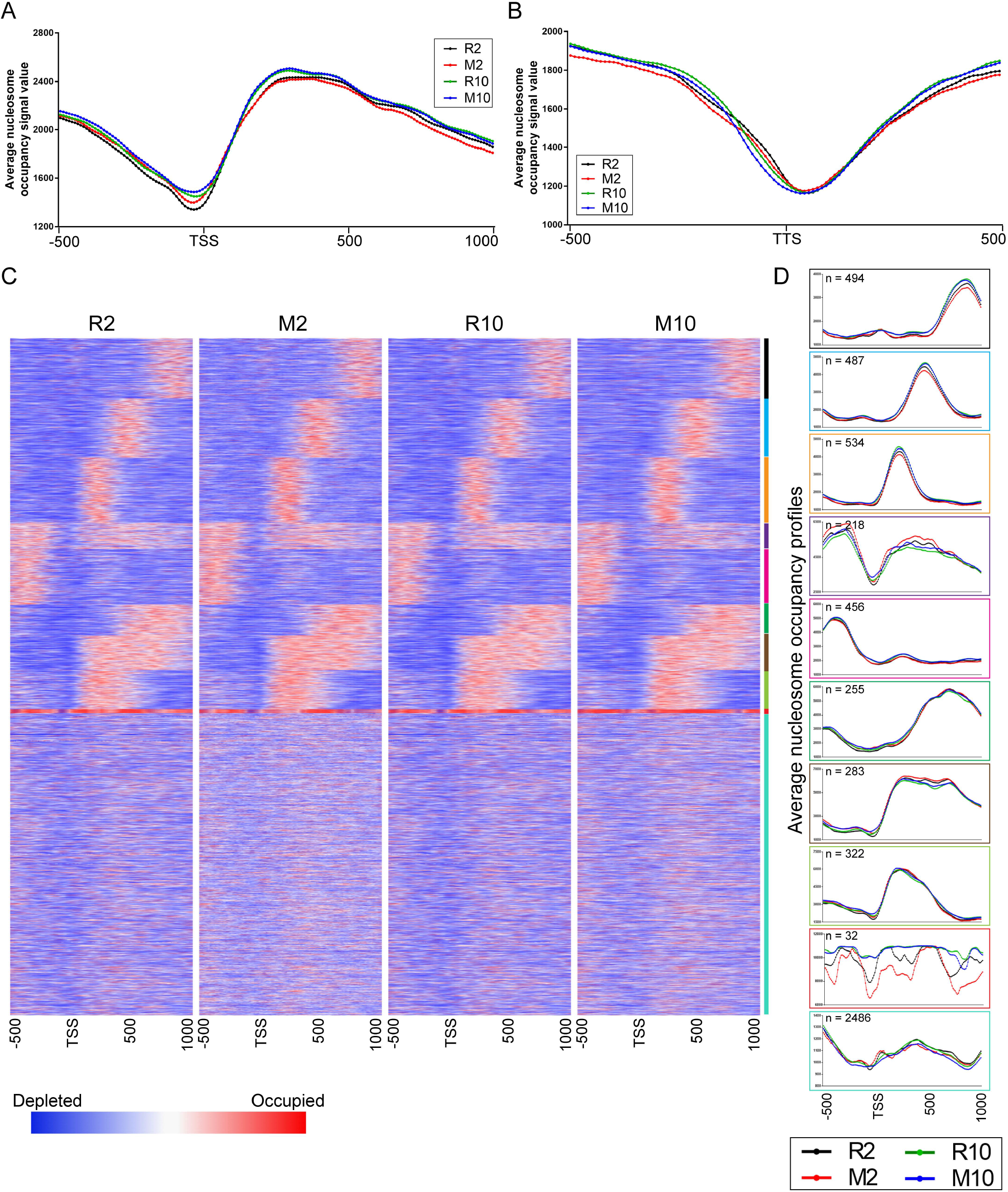
Nucleosome organization in *Cg*. **A and B,** Overlays of average nucleosome occupancy profiles of 5300 *Cg* genes, relative to the transcription start site (TSS) **(A)** and the transcription termination site (TTS) **(B)**. R2: 2 h RPMI-grown, R10: 10 h RPMI-grown, M2: 2 h macrophage-internalized and M10: 10 h macrophage-internalized *Cg* cells. **C,** Heatmaps depicting global nucleosome occupancy profiles around TSS which are clustered into 10 distinct gene clusters. **D,** Overlays of gene-cluster-specific nucleosome occupancy profiles around TSS. The box color corresponds to the color code of each cluster depicted in panel *c*.

**Appendix Figure S2:**
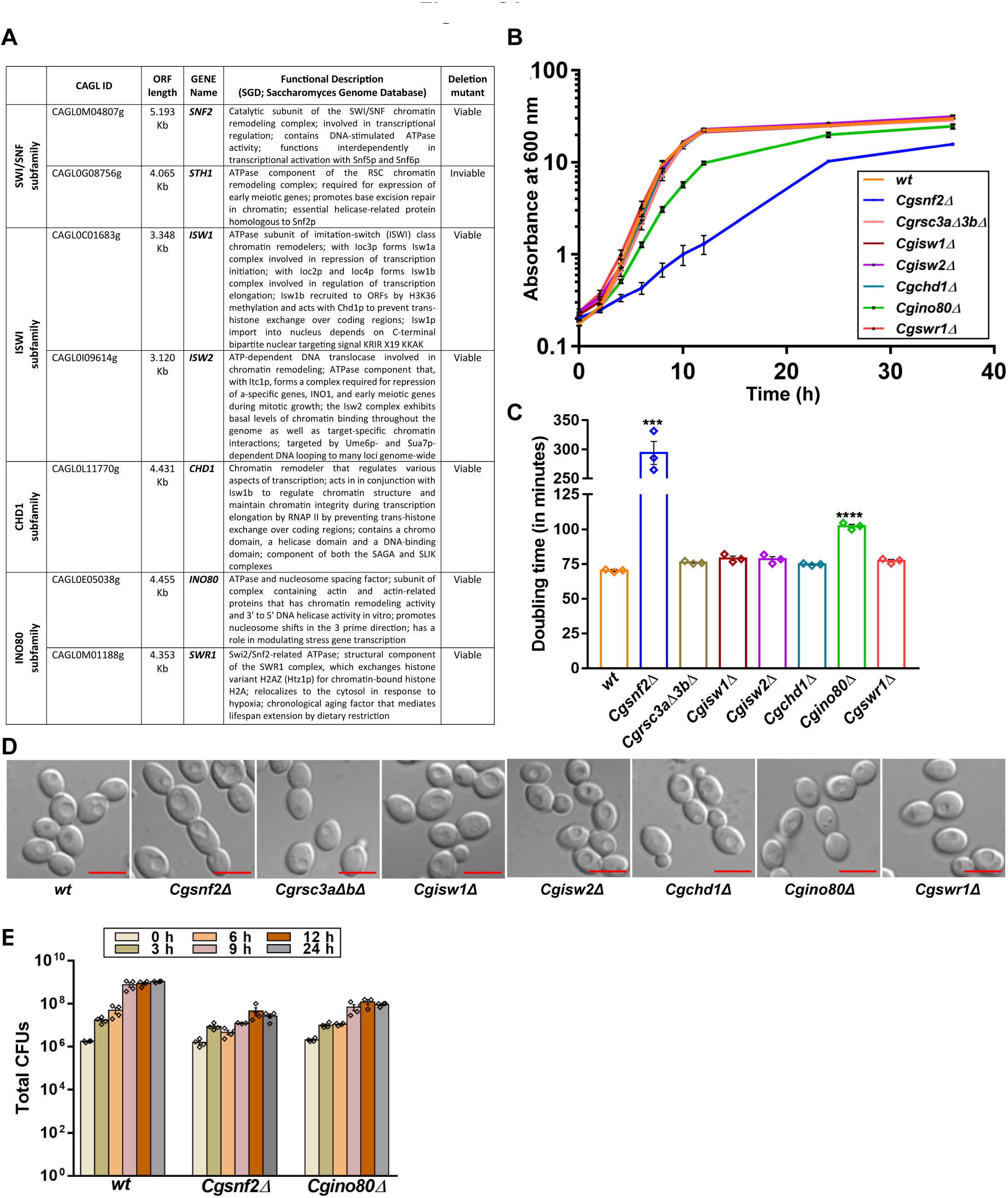
*CgSNF2* deletion leads to growth retardation. **A,** ATPase subunits of seven ATP-dependent chromatin remodelling complexes in *C. glabrata*, as identified by BLAST analysis of their *S. cerevisiae* orthologs. **B,** Time course analysis of *Cg* growth in YPD medium. **C,** Generation time, calculated during the logarithmic-phase (between 2 and 8 h) of the growth curve shown in *b* panel. Data represent mean ± SEM (n = 3). Unpaired two-tailed Student’s *t* test. **D,** Representative confocal images of YPD-grown, log-phase *Cg* cells. Scale bar = 5 µm. **E,** CFU-based growth analysis of *Cg* strains in RPMI medium containing 10% FBS at 37°C, 5% CO_2_. At indicated time points, appropriate dilutions of RPMI-grown cultures were plated on YPD medium. After incubation at 30°C for 2-3 days, colonies were counted and plotted. Data represent mean ± SEM (n = 3-4).

**Appendix Figure S3:**
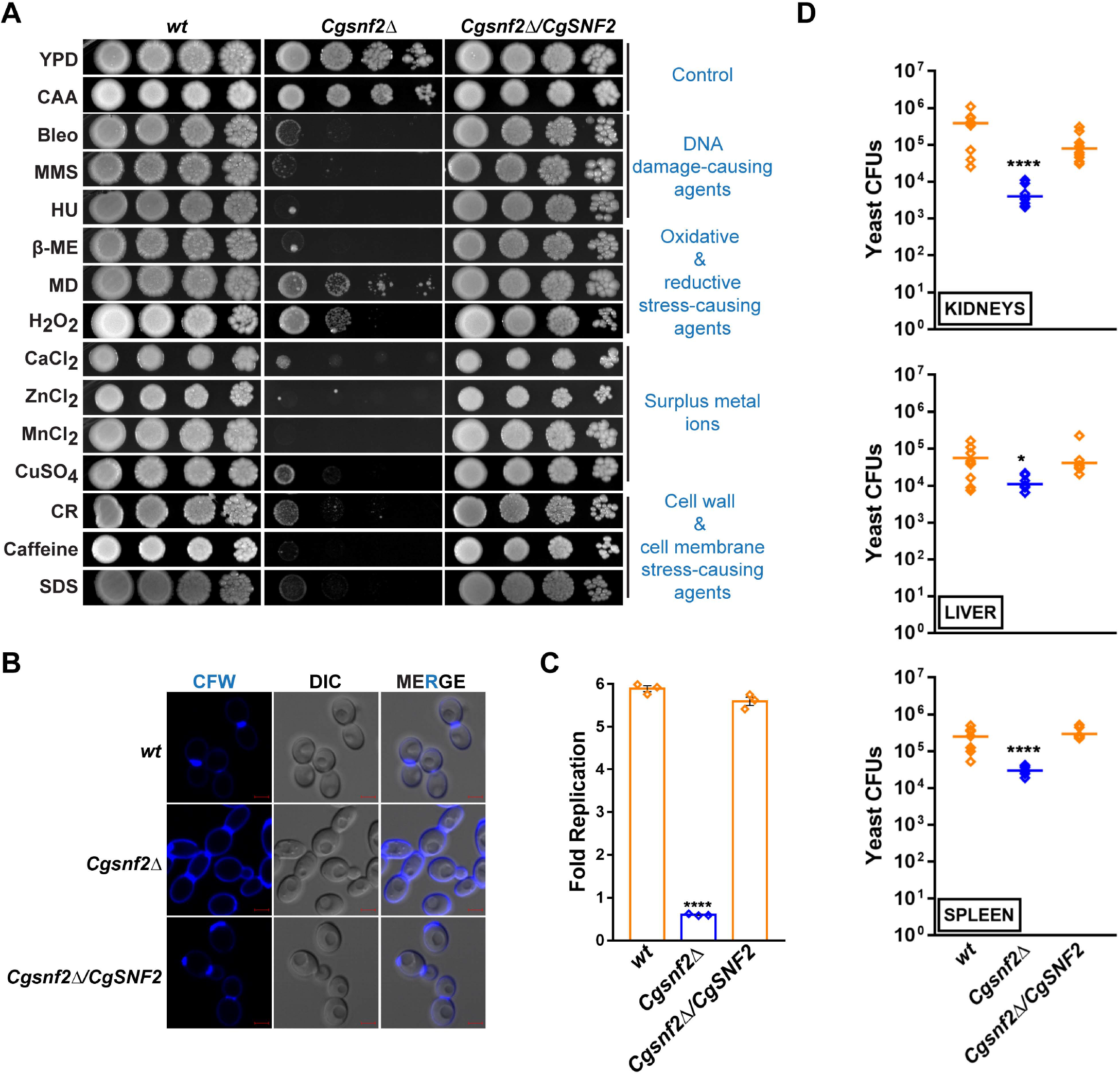
*CgSNF2* complements virulence defect of *Cgsnf2Δ*. **A,** Serial dilution spotting assay showing *Cg* growth. Bleomycin (Bleo), methyl methanesulfonate (MMS), hydroxyurea (HU), β -mercaptoethanol (β-ME), menadione (MD), hydrogen peroxide (H_2_O_2_), calcium chloride (CaCl_2_), zinc chloride (ZnCl_2_), manganese chloride (MnCl_2_), copper sulphate (CuSO_4_), congo red (CR), caffeine and sodium dodecyl sulfate (SDS) were used at 5 μM, 0.03%, 25 mM, 20 mM, 100 µM, 25 mM, 500 mM, 8 mM, 3 mM, 8 mM, 2 mg/ml, 10 mM and 0.05% concentrations, respectively. **B,** Representative confocal images of log-phase CAA medium-grown, calcofluor white-stained *Cg* cells. Scale bar = 2 µm. **C,** CFU-based intracellular survival analysis of *Cg* strains in THP-1 macrophages. Data represent mean ± SEM (n = 3). Unpaired two-tailed Student’s t-test. **D,** CFU-based analysis of organ fungal burden in the murine model of systemic candidiasis. Diamonds and horizontal line represent CFUs recovered from each mouse and geometric CFU mean (n = 8-10) for each organ, respectively. Mann-Whitney test. CFU values plotted for *wt* and *Cgsnf2Δ* strains are the same, as shown in Figure 2D.

**Appendix Figure S4:**
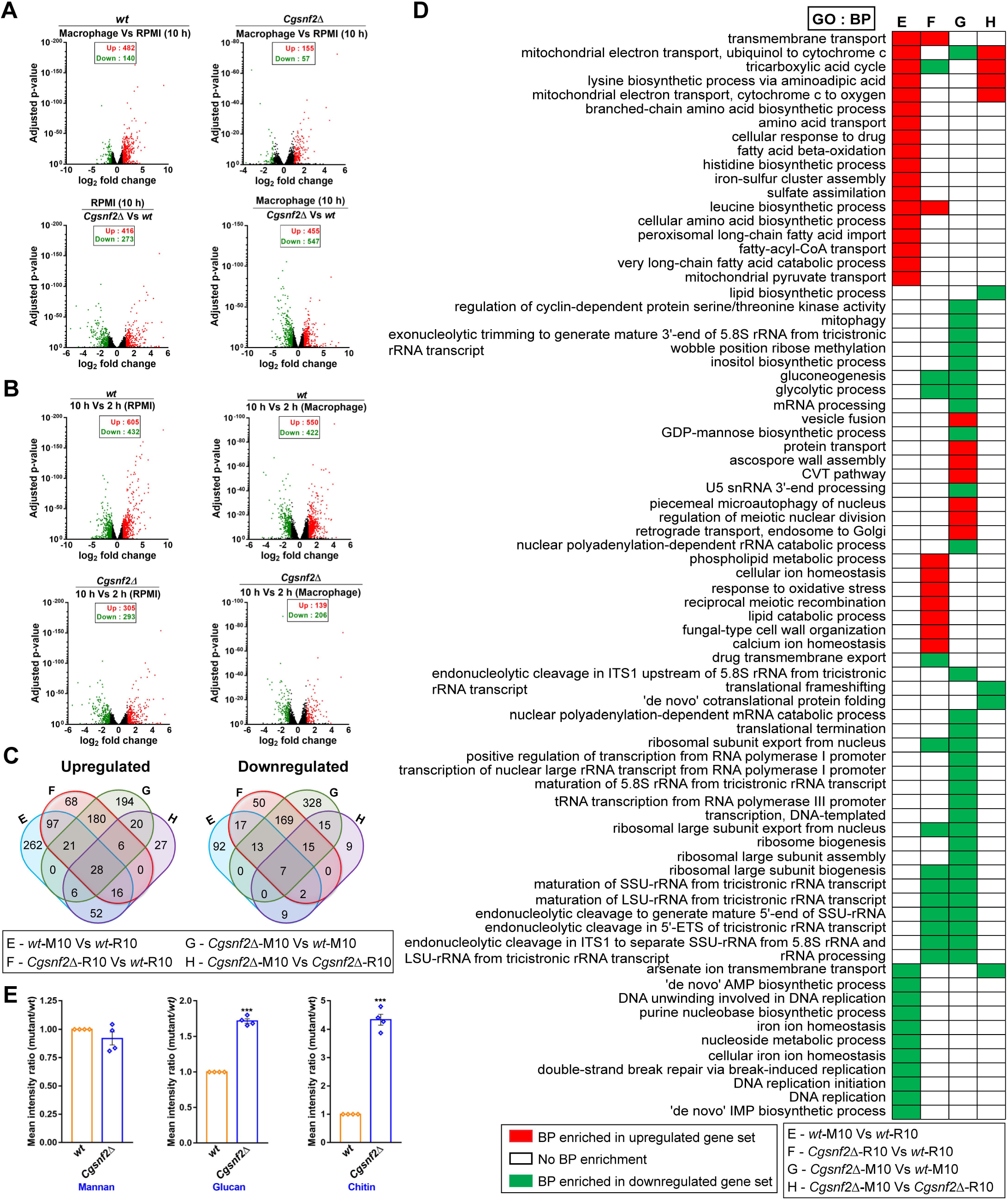
CgSnf2 regulates gene expression. **A and B,** Volcano plots showing mean fold change of differentially-expressed genes (DEGs) and their adjusted p (false discovery rate-corrected) values in 10 h **(A)** and 10 h vs 2 h comparison **(B)**. Red, green and black dots indicate upregulated, downregulated and non-DEGs, respectively. **C,** Venn diagrams showing overlap in DEGs among four compared datasets. **D,** Heatmap showing enriched GO-BP terms in four compared datasets. **E,** Bar graph showing mannan, glucan and chitin amounts in *Cgsnf2Δ*, relative to *wt*. Data represent mean fluorescence intensity ratios ± SEM (n = 4). Paired two-tailed Student t test.

**Appendix Figure S5:**
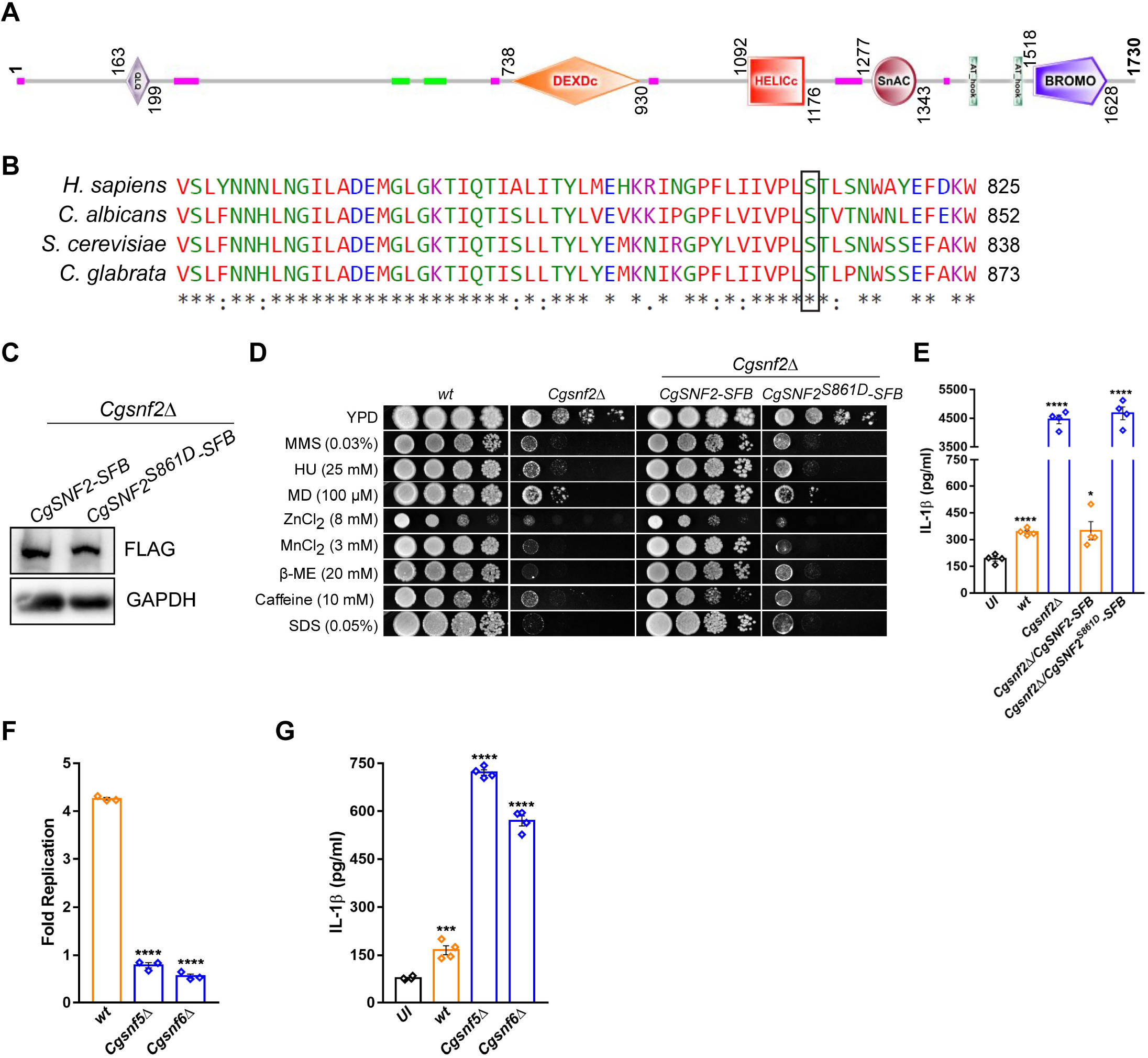
The conserved serine-861 residue in the ATPase domain of CgSnf2 is required for the host immune response suppression. **A,** Schematic representation of different domains in CgSnf2 (CAGL0M04807p), as predicted by the SMART tool (http://smart.embl-heidelberg.de/). QLQ: Gln, Leu, Gln motif; DEXDc: DEAD-like helicases superfamily; HELICc: helicase superfamily c-terminal domain; SnAC: Snf2-ATP coupling, chromatin remodelling complex; AT_hook: DNA binding domain with preference for A/T rich regions and BROMO: bromo domain. Green and pink-colored regions represent coiled-coil and low-complexity regions, respectively. The SnAC domain is involved in binding to histones and required for mobilising nucleosomes. **B,** Multiple amino acid sequence alignment, generated using CLUSTAL-W program, illustrating the conserved serine-861 residue in CgSnf2. **C,** Representative western blot showing similar expression of *wild-type* and serine-to-aspartate-mutated (CgSnf2*^S861D^*) CgSnf2 tagged with the triple epitope SFB [S protein-Flag-Streptavidin-binding peptide] at the C-terminus, as detected using anti-FLAG antibody. **D,** Serial dilution spotting analysis illustrating non-complementation of *Cgsnf2Δ* mutant phenotypes by CgSnf2*^S861D^*. Methyl methanesulfonate (MMS), hydroxyurea (HU), menadione (MD), zinc chloride (ZnCl_2_), manganese chloride (MnCl_2_), β -mercaptoethanol (β-ME), caffeine and sodium dodecyl sulfate (SDS) were used at 0.03%, 25 mM, 100 µM, 8 mM, 3 mM, 20 mM, 10 mM and 0.05% concentrations, respectively. **E,** IL-1β secretion in THP-1 macrophages. Data represent mean ± SEM (n = 4). Unpaired two-tailed Student’s t-test. **F,** *Cg* survival analysis in THP-1 macrophages. Data represent mean ± SEM (n = 3). Unpaired two-tailed Student’s t-test. **G,** IL-1β secretion in THP-1 macrophages. Data represent mean ± SEM (n = 4). Unpaired two-tailed Student’s t-test.

**Appendix Figure S6:**
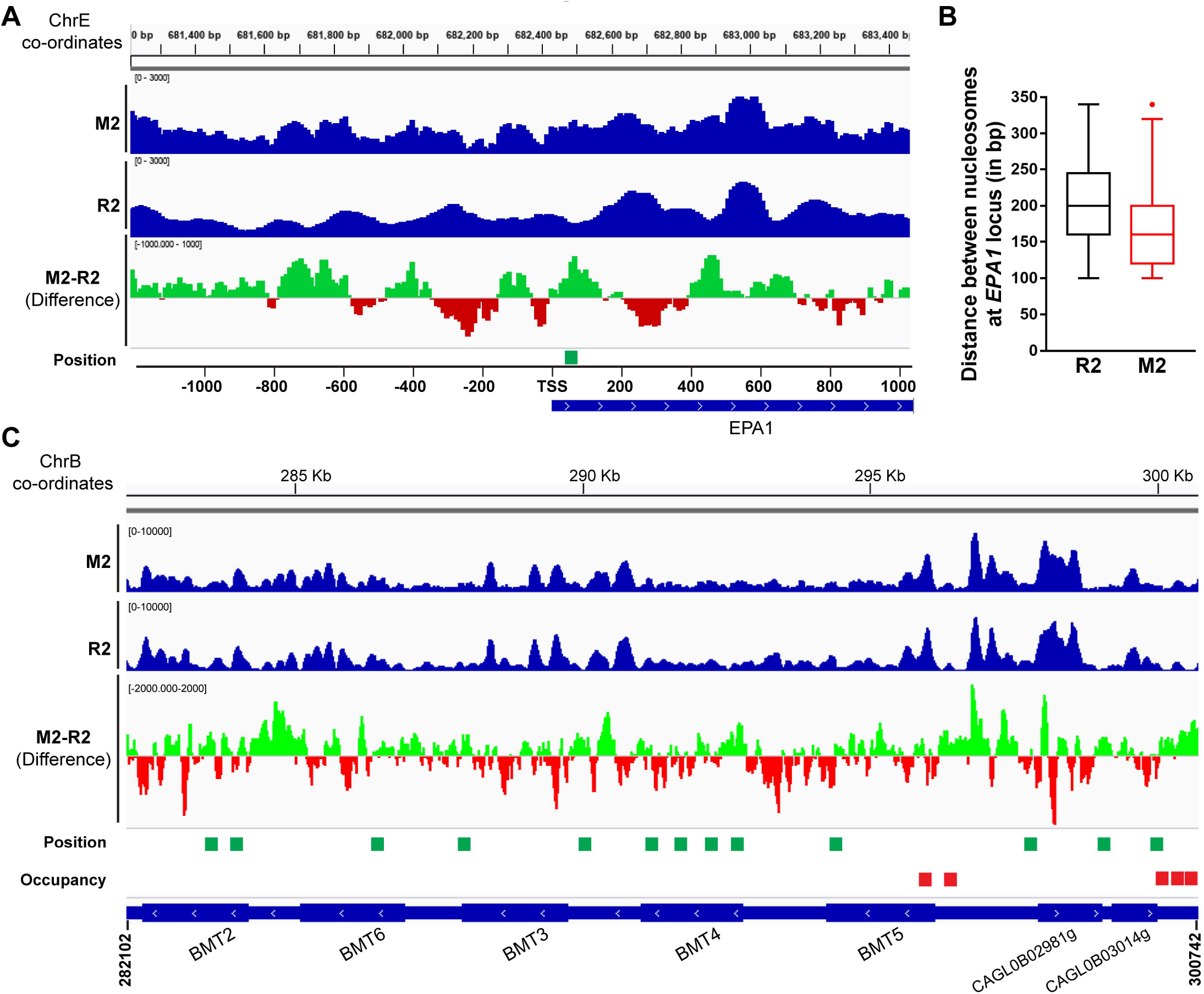
CgSnf2 binds to *EPA1* promoter. **A and C,** Integrative genome viewer (IGV) snapshot of MNase-seq signal at *EPA1* **(A)** and *MT* **(C)** loci in *wild-type* cells grown either in RPMI-medium (R2) or incubated with THP-1 macrophages (M2) for 2 h. Green and red peaks in M2-R2 indicate an increase and a decrease in nucleosome occupancy, respectively, upon macrophage internalization. The scaling factor for the Y-axis is indicated at the top of each IGV track. The arrow indicates the direction of transcription. **B,** Nucleosomes at *EPA1* locus (ChrE: 681312 – 686692 bp) are more compactly arranged in 2 h macrophage-internalized (M2), compared to 2 h RPMI-grown *Cg* (R2). Dynamic nucleosomes are shown underneath both *EPA1* and *CgMT-C* IGV tracks.

**Appendix Table S1 to S10**

**Appendix Table S1: Nucleosome mapping in *C. glabrata*** (***Cg*).**

**Appendix Table S1A:** Total nucleosomes identified in 2 h RPMI-grown *wild-type* (w*t*) cells.

**Appendix Table S1B:** Total nucleosomes identified in 2 h macrophage-internalized *wt* cells.

**Appendix Table S1C:** Total nucleosomes identified in 10 h RPMI-grown *wt* cells.

**Appendix Table S1D**: Total nucleosomes identified in 10 h macrophage-internalized *wt* cells.

**Appendix Table S2: Nucleosome dynamics in *Cg*.**

**Appendix Table S2A:** Nucleosomes showing position shifts in 2 h macrophage-internalized *Cg*, as compared to 2 h RPMI-grown *Cg*.

**Appendix Table S2B:** Nucleosomes showing occupancy changes in 2 h macrophage-internalized *Cg*, as compared to 2 h RPMI-grown *Cg*.

**Appendix Table S2C:** Nucleosomes showing fuzziness changes in 2 h macrophage-internalized *Cg*, as compared to 2 h RPMI-grown *Cg*.

**Appendix Table S2D:** Nucleosomes showing position shifts in 10 h macrophage-internalized *Cg*, as compared to 10 h RPMI-grown *Cg*.

**Appendix Table S2E:** Nucleosomes showing occupancy changes in 10 h macrophage-internalized *Cg*, as compared to 10 h RPMI-grown *Cg*.

**Appendix Table S2F:** Nucleosomes showing fuzziness changes in 10 h macrophage-internalized *Cg*, as compared to 10 h RPMI-grown *Cg*.

**Appendix Table S2G:** Nucleosomes showing position shifts in 10 h macrophage-internalized *Cg*, as compared to 2 h macrophage-internalized *Cg*.

**Appendix Table S2H:** Nucleosomes showing occupancy changes in 10 h macrophage-internalized *Cg*, as compared to 2 h macrophage-internalized *Cg*.

**Appendix Table S2I:** Nucleosomes showing fuzziness changes in 10 h macrophage-internalized *Cg*, as compared to 2 h macrophage-internalized *Cg*.

**Appendix Table S2J:** Nucleosomes showing position shifts in 10 h RPMI-grown *Cg*, as compared to 2 h RPMI-grown *Cg*.

**Appendix Table S2K:** Nucleosomes showing occupancy changes in 10 h RPMI-grown *Cg*, as compared to 2 h RPMI-grown *Cg*.

**Appendix Table S2L:** Nucleosomes showing fuzziness changes in 10 h RPMI-grown *Cg*, as compared to 2 h RPMI-grown *Cg*.

**Appendix Table S3: Nucleosome dynamics at gene promoters in *Cg*.**

**Appendix Table S3A:** Genes with dynamic nucleosomes at their promoter regions in 2 h macrophage-internalized *Cg*, as compared to 2 h RPMI-grown *Cg*.

**Appendix Table S3B:** Genes with dynamic nucleosomes at their promoter regions in 10 h macrophage-internalized *Cg*, as compared to 10 h RPMI-grown *Cg*.

**Appendix Table S3C:** Genes with dynamic nucleosomes at their promoter regions in 10 h macrophage-internalized *Cg*, as compared to 2 h macrophage-internalized *Cg*.

**Appendix Table S3D:** Genes with dynamic nucleosomes at their promoter regions in 10 h RPMI-grown *Cg*, as compared to 2 h RPMI-grown *Cg*.

**Appendix Table S4: Gene Ontology (GO) enrichment analysis for *Cg* genes that contain dynamic nucleosomes at their promoters.**

**Appendix Table S4A:** Enriched GO terms in 2 h macrophage-internalized *Cg*, as compared to 2 h RPMI-grown *Cg*.

**Appendix Table S4B:** Enriched GO terms in 10 h macrophage-internalized *Cg*, as compared to 10 h RPMI-grown *Cg*.

**Appendix Table S4C:** Enriched GO terms in 10 h macrophage-internalized *Cg*, as compared to 2 h macrophage-internalized *Cg*.

**Appendix Table S4D:** Enriched GO terms in 10 h RPMI-grown *Cg*, as compared to 2 h RPMI-grown *Cg*.

**Appendix Table S5: Differential gene expression in response to macrophage internalization and *CgSNF2* deletion.**

**Appendix Table S5A**: Differentially-expressed genes (DEGs) in 2 h macrophage-internalized *wild-type* (*wt*), as compared to 2 h RPMI-grown *wt*.

**Appendix Table S5B:** DEGs in 2 h RPMI-grown *Cgsnf2*Δ, as compared to 2 h RPMI-grown *wt*.

**Appendix Table S5C:** DEGs in 2 h macrophage-internalized *Cgsnf2*Δ, as compared to 2 h RPMI-grown *Cgsnf2*Δ.

**Appendix Table S5D:** DEGs in 2 h macrophage-internalized *Cgsnf2*Δ, as compared to 2 h macrophage-internalized *wt*.

**Appendix Table S5E:** DEGs in 10 h macrophage-internalized *wt*, as compared to 10 h RPMI-grown *wt*.

**Appendix Table S5F:** DEGs in 10 h RPMI-grown *Cgsnf2*Δ, as compared to 10 h RPMI-grown *wt*.

**Appendix Table S5G:** DEGs in 10 h macrophage-internalized *Cgsnf2*Δ, as compared to 10 h RPMI-grown *Cgsnf2*Δ.

**Appendix Table S5H:** DEGs in 10 h macrophage-internalized *Cgsnf2*Δ, as compared to 10 h macrophage-internalized *wt*.

**Appendix Table S5I:** DEGs in the 10 h RPMI-grown *wt*, as compared to 2 h RPMI-grown *wt*.

**Appendix Table S5J:** DEGs in 10 h macrophage-internalized *wt*, as compared to 2 h macrophage-internalized *wt*.

**Appendix Table S5K:** DEGs in 10 h RPMI-grown *Cgsnf2*Δ, as compared to 2 h RPMI-grown *Cgsnf2*Δ.

**Appendix Table S5L:** DEGs in 10 h macrophage-internalized *Cgsnf2*Δ, as compared to 2 h macrophage-internalized *Cgsnf2*Δ.

**Appendix Table S6: Gene Ontology (GO) enrichment analysis for differentially-expressed genes.**

**Appendix Table S6A:** Enriched GO terms in 2 h macrophage-internalized *wt*, as compared to 2 h RPMI-grown *wt*.

**Appendix Table S6B:** Enriched GO terms in 2 h RPMI-grown *Cgsnf2*Δ, as compared to 2 h RPMI-grown *wt*.

**Appendix Table S6C:** Enriched GO terms in 2 h macrophage-internalized *Cgsnf2*Δ, as compared to 2 h RPMI-grown *Cgsnf2*Δ.

**Appendix Table S6D:** Enriched GO terms in 2 h macrophage-internalized *Cgsnf2*Δ, as compared to 2 h macrophage-internalized *wt*.

**Appendix Table S6E:** Enriched GO terms in 10 h macrophage-internalized *wt*, as compared to 10 h RPMI-grown *wt*.

**Appendix Table S6F:** Enriched GO terms in 10 h RPMI-grown *Cgsnf2*Δ, as compared to 10 h RPMI-grown *wt* cells.

**Appendix Table S6G:** Enriched GO terms in 10 h macrophage-internalized *Cgsnf2*Δ, as compared to 10 h RPMI-grown *Cgsnf2*Δ.

**Appendix Table S6H:** Enriched GO terms in 10 h macrophage-internalized *Cgsnf2*Δ, as compared to 10 h macrophage-internalized *wt*.

**Appendix Appendix Table S6I:** Enriched GO terms in 10 h RPMI-grown *wt*, as compared to 2 h RPMI-grown *wt*.

**Appendix Table S6J:** Enriched GO terms in 10 h macrophage-internalized *wt*, as compared to 2 h macrophage-internalized *wt*.

**Appendix Table S6K:** Enriched GO terms DEGs in 10 h RPMI-grown *Cgsnf2*Δ, as compared to 2 h RPMI-grown *Cgsnf2*Δ.

**Appendix Table S6L:** Enriched GO terms in 10 h macrophage-internalized *Cgsnf2*Δ, as compared to 2 h macrophage-internalized *Cgsnf2*Δ.

**Appendix Table S7: List of *C. glabrata* strains used in the study.**

**Appendix Table S8: List of plasmids used in the study.**

**Appendix Table S9: List of primers used in the study.**

**Appendix Table S10: List of antibodies and inhibitors used in the study.**

